# D-amino acid substitution and cyclisation enhance the stability and antimicrobial activity of arginine-rich peptides

**DOI:** 10.1101/2025.06.17.660067

**Authors:** Bruno Mendes, Valeria Castelletto, Ian W. Hamley, Glyn Barrett

## Abstract

Cationic peptides rich in arginine are potent antimicrobials but face challenges due to enzymatic degradation and potential host toxicity. We screened eight linear arginine-rich peptides for antimicrobial and haemolytic activity, identifying R4F4 as the most active. However, its efficacy decreased in the presence of human serum and trypsin. To enhance stability, we developed three R4F4 derivatives—D-R4F4, cyclic R4F4, and R4F4-C16—plus one R4-based peptide (R4-C16). Cyclisation and D-amino acid substitution significantly improved protease resistance and antimicrobial activity. Using fluorescence imaging, microscopy, and RNA sequencing, we revealed a complex mechanism involving membrane disruption, oxidative stress modulation, and transcriptomic changes in bacterial metabolic pathways. The R4F4-based peptides also displayed strong antibiofilm activity and good cytocompatibility. Our findings highlight the benefits of peptide optimisation—particularly D-amino acid incorporation and cyclisation—in improving stability and function. This work informs future design of antimicrobial peptides and deepens understanding of their multifaceted biological effects.

## 1. INTRODUCTION

Antimicrobial resistance remains a pressing global health issue that continues to escalate as bacteria, through natural selection and mutation, develop sophisticated mechanisms to evade the toxic effects of conventional antibiotics [1]. The dearth in development of novel, effective antibiotics leads to higher rates of mortality and morbidity, disproportionately affecting less developed parts of the world. The discovery and introduction of new anti-infective drugs into the market is lengthy, expensive, and most often with limited success. These factors collectively underscore the growing urgency to intensify efforts toward discovering and developing alternative and effective therapeutic agents.

The traditional screening of naturally derived compounds and the use of bioinformatics and artificial intelligence (AI) for mining large genomes and proteomes have led to the discovery of many natural peptides with antimicrobial properties [2,3]. Bioactive peptides are traditionally characterised by a high content of positively charged and hydrophobic amino acids, which enable them to disrupt bacterial membranes and induce cell death [3,4]. The differential distribution of phospholipids in eukaryotic and prokaryotic cells affects the initial electrostatic and hydrophobic interactions of peptides, conferring a degree of selectivity that supports their potential for drug development and safety in clinical settings. Consequently, many studies have focused on designing, developing and/or isolating peptide-based antimicrobials encoding lysine or arginine residues strategically integrated with regions with hydrophobic properties [5–7]. Our group and other research teams, for instance, have discovered a series of short amphiphilic peptides with enhanced antibacterial activity, effectively combining positively charged and hydrophobic amino acids [8–13]. Although this strategy has been promising in identifying potent antimicrobial agents, their toxicity and low stability in serum conditions have hindered clinical translation and further progress [14,15]. In the presence of serum, peptides can aggregate through hydrophobic interactions, leading to a reduction in their effective concentration and bioavailability. Furthermore, linear peptides often adopt flexible conformations, making them even more susceptible to proteolysis. On the other hand, an imbalance of positive charges and hydrophobic properties in the peptide sequence can shift its preference toward human cells, enhancing interactions that may compromise the integrity and functionality of red blood cells and other cell types.

Successful cases of peptide-based drug development have been rooted in modified analogues with lower toxicity and higher stability [15,16]. Chemical modifications, such as cyclisation, PEGylation, lipidation and incorporation of D-amino acids have played significant roles in the development and positive activity and clinical applicability of peptide-based antibiotics [17,18]. Classic examples of now routinely used peptide-based antimicrobials include Daptomycin and Polymyxin B (both cyclic lipopeptides); and Oritavancin (a glycopeptide) [19,20]. These commercially available lipopeptides were modified through the glycosylation, the formation of cyclic structures, and the incorporation of D-amino acids; structural features that improve stability and efficacy [21,22]. With this in mind, our study explored the computational design and experimental screening of both the antimicrobial and haemolytic activities of eight linear arginine-rich peptides. We also designed four chemically modified peptides to investigate the benefits of lipidation, D-amino acid substitution and cyclisation in the activity and stability of novel peptide-based antimicrobials. We also explored the antibacterial mechanism of action, employing microscopic-, fluorescence-, and transcriptomics-based approaches to better understand the diverse range of events induced and modulated through exposure to tested peptides.

## 2. MATERIALS AND METHODS

### 2.1. Computational screening of arginine-rich peptides

Eight arginine-rich peptides were designed for both computational and experimental assessments, focusing on their antimicrobial properties and toxicity (**Supplementary Table S1**). The amphiphilic sequences combine arginine and hydrophobic amino acids, such as alanine, phenylalanine or valine. The design was initially guided by our previous research with cationic peptides and other studies exploring the frequency and distribution of amino acids in peptide-based antimicrobial agents [5,10,23,24]. We used online, open-source predictors to screen the potential antimicrobial activity (AMPfun and Antimicrobial Peptide Scanner vr.2), haemolytic effects (HemoPI), and toxicity (ToxinPred) based on the primary structure and physicochemical properties of the designed peptides.

### 2.2. Synthesis, purification and identification of peptides

All peptides used in this study were synthesised using the Fmoc-strategy and purchased from Peptide Protein Research Ltd. (Hampshire, UK). Based on our experimental screening, four modified peptides were also purchased and included in our analysis (**Supplementary Table S2**). Three were inspired by the R4F4 sequence: R4F4-C16, a R4F4 peptide with C16 palmitoyl chain at N-terminus; D-R4F4, which has the same sequence of R4F4, but with alternated L and D amino acids; and CP-R4F4, presenting a cyclic structure. The last one was R4-C16, which is also an arginine-rich lipopeptide. The purity level was estimated by high-performance liquid chromatography (HPLC), while the molecular mass of the peptides was confirmed by mass spectrometry (MS), as described in our previous research [5]. All synthetic products were confirmed to have a purity at or exceeding 95% (**Supplementary Figure 1**).

### 2.3. Antibacterial assessment

The minimal inhibitory concentration (MIC) of arginine-based peptides was assessed using the microbroth dilution approach [19]. Our *in vitro* screening included a panel of Gram-negative bacteria, *Escherichia coli* (ATCC 25922 and enterohemorrhagic O157:H7) and *Pseudomonas aeruginosa* (PA01 and the drug-resistant strain NCTC 13437), and Gram-positive bacteria (*Staphylococcus aureus* ATCC 12600). 5 mL overnight cultures in Lysogeny broth, prepared from single isolated colonies, were diluted to 5 × 10^6^ CFU mL^−1^ mid-log phase in Muller Hinton (MH) broth. Next, peptide concentrations (0 to 1 mM) were serially diluted two-fold in 96-well plates and incubated overnight at 37 °C with tested bacterial strains. In each case, the MIC was determined as the lowest peptide concentration at which no detectable bacterial growth was observed employing an absorbance-based assay using a Tecan Spark® multimode plate reader. Subsequently, 10 µL aliquots from each peptide dilution were transferred to MH agar (1.5%) plates to determine the minimum bactericidal concentration (MBC) which was assessed by establishing at which concentration no visible growth of bacteria (assessed via colony formation) on the plate was observed.

### 2.4. Impact of arginine-rich peptides on membrane integrity of bacteria

#### 2.4.1. Outer membrane damage

The potential membrane disruption caused by arginine-based peptides was evaluated using the NPN (N-phenyl-1-naphthylamine) fluorescent probe in a hydrophobic environment. Overnight *E. coli* cultures were adjusted to an OD_600_ of 0.5 in 5 mM HEPES buffer (pH 7.4) with 5 mM glucose and transferred to a black 96-well microtiter plate. Afterwards, 10 µM NPN was added and incubated for 15 min at 37 °C. Following this incubation, either a 2× MIC or 1 mM peptide concentration was incubated, and the fluorescence intensity was recorded at an excitation λ = 350 nm, emission λ = 420 nm using a Tecan Spark® plate reader for 1h at 37 °C.

#### 2.4.2. Plasma membrane depolarisation

The dye 3,3′-dipropylthiadicarbocyanine iodide DiSC3(5) was employed to evaluate perturbation of the bacterial membrane by our designed arginine-rich peptides. For this, a black 96-well microtiter plate containing 150 µL *E. coli* cells (OD_600_= 0.5) in MH broth per well was incubated with 0.5 µM DiSC3(5) in 5 mM HEPES buffer (pH 7.4, supplemented with 5 mM glucose and 0.1 M KCl) for 15 min to allow for fluorescence quenching. Then, 2× MIC or 1 mM peptide was added, and changes in fluorescence were monitored in real-time (λex = 622 nm, λem = 670 nm) using a microplate reader.

#### 2.4.3. Cell membrane integrity

Propidium iodide (PI) uptake was used to assess the permeabilisation of the bacterial membrane upon incubation with peptides. *E. coli* and *S. aureus* cells, adjusted to an OD_600_ of 0.5, were placed into a 96-well plate containing 5 mM HEPES buffer (pH 7.4, supplemented with 5 mM glucose) and incubated with 5 µg mL^-1^ PI for 15 min. Afterwards, different peptide concentrations were added, and PI uptake was rapidly measured every 10 min over a 1h period using a Tecan Spark® microplate reader (λex = 515 nm, λem = 620 nm).

#### 2.4.4. Bacterial visualisation using Atomic Force Microscopy (AFM)

Changes in bacterial morphology were analysed using AFM. *E. coli* (5 × 10⁶ CFU mL^-1^, mid-log phase) cells were incubated with two-fold MIC peptides in MH broth for 3h at 37°C. The cells were harvested by resuspension in PBS and fixed overnight with 2.5% glutaraldehyde. After fixation, bacterial cells were washed 3 times in PBS and transferred to 1 cm² pieces of muscovite mica on metal stubs using double-sided tape. AFM imaging was performed using an Asylum Research Cypher S AFM microscope in air tapping mode, with a 30 µm scan size, 512 points line^-1^, and a 0.75 Hz scan rate.

### 2.5. Gene expression profiles of *E. coli* in response to peptide exposure

#### 2.5.1. RNA isolation, library construction, sequencing, and bioinformatics analysis

A comprehensive transcriptome analysis was conducted to gain insights into the mechanism of action of the most active antimicrobial peptide. *E. coli* overnight cultures were adjusted to an OD600 of 0.5 and incubated with the MIC of D-R4F4 in MH broth for 24 h at 37 °C. PBS groups were also included as negative control. After incubation, samples were centrifuged three times at 4000 rpm for 10 min at 4 °C with PBS. Total RNA was extracted from approximately 15 μg of cell pellet using TRIzol, following the manufacturer’s recommendations. RNA quality and quantity were assessed with an Agilent 2100/5400 Bioanalyzer, while concentration was measured using the Qubit™ RNA Assay Kit. Ribosomal RNA was removed using the Ribo-Zero kit to enrich mRNA.

The enriched mRNA was fragmented using an ultrasonicator, and first-strand cDNA was synthesised using random hexamer primers and M-MuLV Reverse Transcriptase (RNase H–). For second-strand cDNA synthesis, dUTPs were incorporated instead of dTTPs to maintain strand specificity. The cDNA fragments underwent end-repair, A-tailing, and adapter ligation, followed by size selection and PCR amplification to enrich adapter-ligated fragments.

The final libraries were purified using the AMPure XP system and assessed for quality, concentration, and fragment size distribution using Qubit, real-time PCR, and an Agilent Bioanalyzer 2100. Libraries were generated to their effective concentration and sequencing according to the requirements before being sequenced on an Illumina HiSeq platform by Novogene (UK) Ltd.

#### 2.5.2. Data Processing and Gene Expression Quantification

Raw sequencing reads were processed using fastp, removing adapter sequences, low-quality reads (Q < 20), and short reads (<50 bp). Clean reads were assessed for quality using Q20, Q30, and GC content. Genome mapping and sequence filtering were performed using Bowtie2 (v2.5.4). Gene expression levels were quantified with featureCounts (v2.0.6), and expression values were normalised as Fragments Per Kilobase of transcript per Million mapped reads (FPKM), accounting for gene length and sequencing depth.

#### 2.5.3. Functional and Pathway Analysis

Gene Ontology (GO) enrichment analysis was conducted using clusterProfiler (v4.8.1) with gene length bias correction, considering GO terms with a corrected P-value < 0.05 as significantly enriched. Kyoto Encyclopedia of Genes and Genomes (KEGG) pathway analysis was also performed with clusterProfiler to investigate high-level biological functions. Additional pathway analyses were conducted using Ingenuity Pathway Analysis (IPA, QIAGEN) and Metacore (Clarivate Analytics). GO enrichment validation was performed using GORILLA and GENEONTOLOGY tools. STRING DATABASE was utilised to construct protein-protein interaction (PPI) networks.

### 2.6. Measurement of intracellular reactive oxygen species (ROS) induced by arginine-rich peptides

Total ROS was measured using an oxidation-sensitive fluorescent dye, 2′,7′-dichlorodihydrofluorescein diacetate (DCFH-DA). For this assay, *E. coli* and *S. aureus* cultures were adjusted to an OD_600_ of 0.3 in MH broth and exposed to peptides at MIC, 1/2× MIC, and 1/4× MIC, or to positive control (5 mM H_2_O_2_) for 4 h at 37 °C. Cells were centrifuged at 4000 rpm for 10 min at 4 °C and resuspended in PBS containing 10 µM DCFH-DA. The suspension was incubated under the same conditions for 30 min. Finally, cells were washed twice with PBS, and 150 µL aliquots was transferred to 96-well plates. Oxidised DCF products were detected using a Tecan Spark® microplate reader with excitation at 485 nm and emission at 530 nm. Additionally, 10 µL of *E. coli* cells incubated with D-R4F4 at MIC, 1/2× MIC, and 1/4× MIC were collected for imaging using an All-in-One Fluorescence Microscope (Keyence, Japan).

### 2.7. Antibiofilm properties

#### 2.7.1. Effect of arginine-rich peptides on early stages of biofilm development

Arginine-rich peptides were evaluated for their ability to inhibit biofilm formation of *P. aeruginosa* (PA01) cells. To assess this, the procedure described in section 2.3 for determining the MIC was adapted using M63 media (supplemented with 1 mM MgSO_4_ and 23 µM L-arginine) [25]. After 24 h of exposure to peptides (0 to 1 mM), flat-bottom 96-well plates were washed twice with PBS to remove any non-adherent cells. The remaining biofilm was stained with 0.1% crystal violet (CV) for 30 min at room temperature. Subsequently, the plates were washed three times to remove excess dye and left to dry overnight at room temperature. The following day, stained biofilms were solubilised using 30% glacial acetic acid, and absorbance was measured at 550 nm using a microplate reader. Biofilm inhibition was expressed as a percentage relative to control bacterial cells treated with PBS.

#### 2.7.2. Effect of arginine-rich peptides on pre-established biofilms

To establish mature *P. aeruginosa* (PAO1) biofilms, an overnight culture was diluted to an OD600 of 0.01 in 120 µL of M63 medium (supplemented with 1 mM MgSO₄ and 23 µM L-arginine) and incubated in flat-bottom 96-well plates at 37 °C for 48 h. Afterwards, individual wells containing biofilm were rinsed three times with PBS, and 150 µL MH broth was added with either 0.5× and 2× MIC peptides. 100 µg mL^-1^ resazurin was added to measure the biofilm metabolic activity for 18h at 37 °C using a Tecan Spark® microplate reader (λex= 520 nm/λem= 590 nm). CFU counts were established by plating aliquots of a dilution series onto MH agar followed by overnight incubation at 37°C.

Three-dimensional images of biofilm communities, with and without peptide exposure, were captured using an All-in-One Fluorescence Microscope (Keyence, Japan). Pre-established biofilms in a 96-well plate were incubated with 2× MIC peptides for 3 h at 37 °C. Dual staining was performed with 5 µL of SYTO 9/PI for 30 min. Biofilm thickness was assessed through 2.0 μm optical cross-sections with 50 μm pinholes.

### 2.8. Impact of serum and protease on the stability and activity of peptides

The antimicrobial activity and structural stability of arginine-rich peptides were evaluated in the presence of human serum (20% and 50%) and 5 μM Sequencing Grade Modified Trypsin (SGMT, Promega). Briefly, fresh human blood was collected in EDTA tubes and allowed to clot at room temperature for 1 h. Serum was then collected following centrifugation at 2000 RPM for 10 min at 4°C. Peptides were incubated with 20%, 50% human serum and 5 μM SGMT in 50 mM Tris-HCl buffer (1 mM CaCl₂, pH 7.6) for 24 h at 37°C. After the incubation period, the solutions were further incubated with *E. coli* (ATCC 25922 and O157:H7), *P. aeruginosa* (PA01, NCTC13437), and *S. aureus* (ATCC12600), and the MIC was assessed as described in section 2.3. Additionally, the samples were analysed using electrospray ionisation mass spectrometry to monitor structural changes based on the molecular weight of the peptides.

### 2.9. *In vitro* haemolysis and cytotoxicity screening

The toxicity screening of arginine-rich peptides was conducted in two cell types: human red blood cells (hRBCs) and L929 fibroblasts. The peptide-induced lysis of hRBCs was estimated based on the release of haemoglobin into solution, following a previously established protocol with slight modifications [26]. Briefly, fresh human blood was collected from a healthy donor. To isolate hRBCs from plasma, the blood was centrifuged at 2000 RPM for 10 min at 4°C, and the cells were carefully resuspended in 0.5% suspension buffer. After, they were added to a 96-well plate containing various concentrations of peptides diluted in PBS. After incubation for 1 h at 37 °C, cells were spun at 2000 RPM for 10 min at 4°C to pellet cell debris and the supernatant collected and measured at OD_414_. The data was calculated as the % haemolysis relative to the positive control (cells treated with 1% Triton X-100). The toxicity screening using L929 fibroblasts was based on the metabolic activity of cells using an MTS colourimetric assay. Initially, the adherent cells were cultured in T75 flasks in Dulbecco’s modified Eagle’s (DMEM) supplemented with 2 mM glutamine, 1% penicillin/streptomycin, and 10% FBS. A total of 10^5^ cells were seeded into individual wells of a 96-well plate, and different peptide concentrations ranging from 0 to 2 mM were added. After 24 h at 37°C and 5% CO₂. 0.3 mg mL^-1^ MTS was added to each well and incubated for 3 h under the same conditions. The absorbance was measured at OD_490_ using a Tecan Spark multimode plate reader. The quantification of viable cells was expressed as a % and determined by comparing peptide-treated groups with control cells grown in their absence. All experiments were performed in triplicate and repeated 2–3 times independently.

### 2.10. Statistical analysis

Statistical analysis was carried out using one-way analysis of variance (ANOVA) with Tukey’s Honest Significant Difference (HSD) post hoc test in GraphPad Prism 9.5. Statistical significance was considered at a p-value < 0.001.

## 3. RESULTS

### 3.1. Computational analyses reveal arginine-rich peptides as promising antimicrobial candidates with low predicted toxicity

Screening the sequences of arginine-rich peptides using webservers AMPFun suggested that all eight linear arginine-rich peptides possessed antimicrobial properties. However, the Antimicrobial Peptide Scanner v2 predicted R4A4, R4V4, and RRARSAVAS to be of limited antimicrobial efficacy, thus contradicting from other prediction tools. Regarding toxicity, HemoPi tools suggested that all peptides are non-haemolytic, with a low tendency to lyse hRBCs (**Table 1**). Similarly, the *in silico* analysis using ToxinPred indicates that arginine-rich peptides potentially lack toxicity to human cells. Initial screening indicates that the designed arginine-rich peptides are likely to be effective against bacteria, with minimal predicted off-target effects based on haemolysis and toxicity assessments..

**Table 1.**
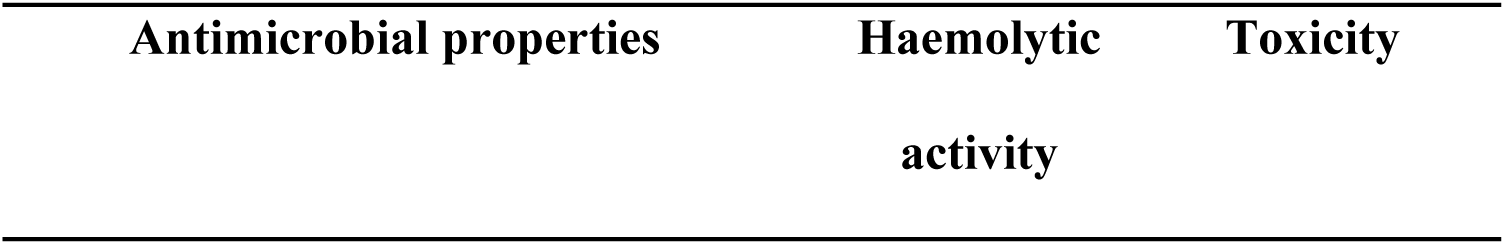
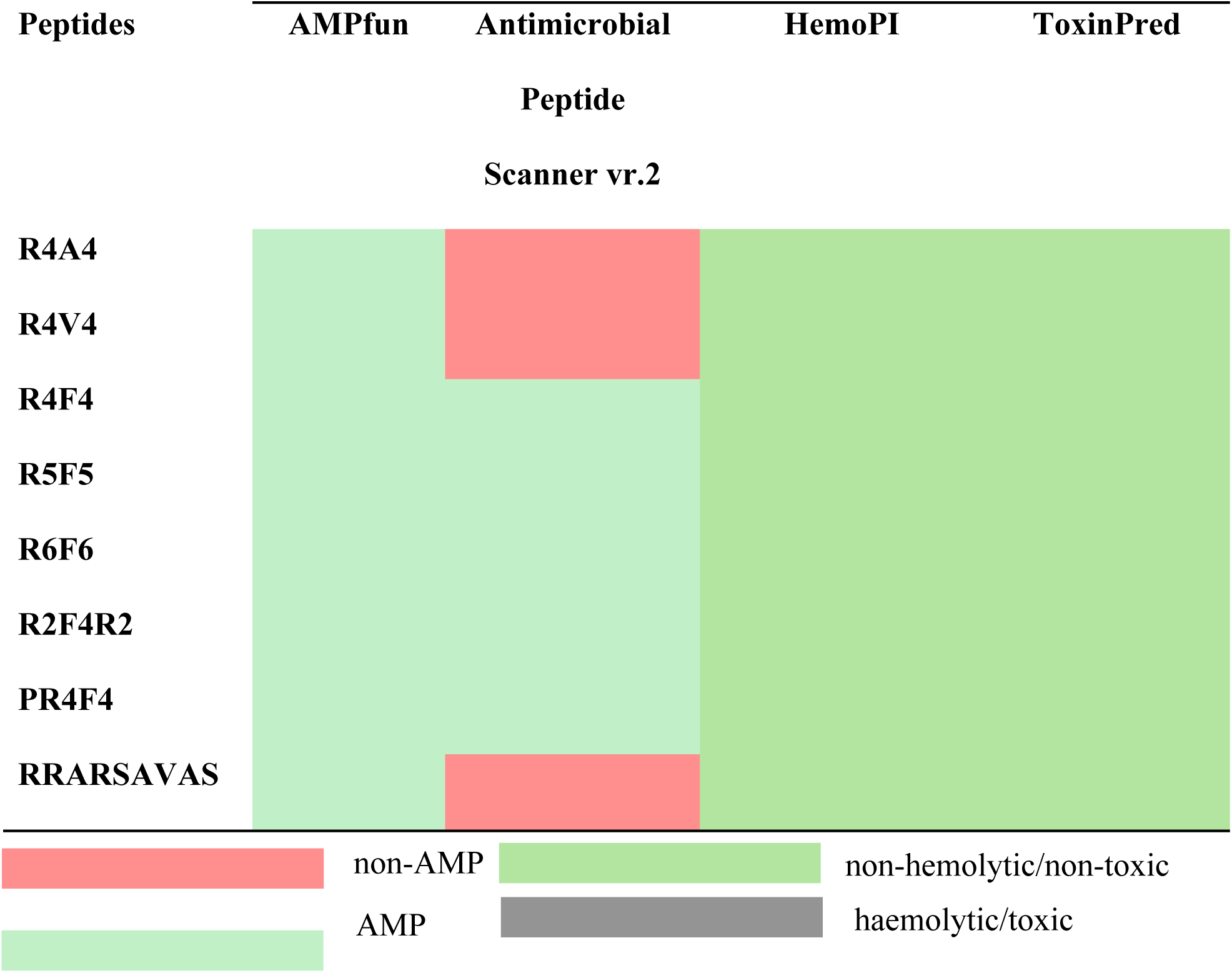
Potential antimicrobial action and non-toxicity of arginine-rich designed peptides. *In silico* screening predictions of antimicrobial, haemolytic activity, and toxicity of linear arginine-rich peptides were conducted using their primary sequences and open sources web servers. IA approaches: AMPfun (http://fdblab.csie.ncu.edu.tw/AMPfun/run.html), Antimicrobial Peptide Scanner vr.2 (https://www.dveltri.com/ascan/v2/ ), HemoPI (https://webs.iiitd.edu.in/raghava/hemopi/batch.php); ToxinPred (https://webs.iiitd.edu.in/raghava/toxinpred/index.html)

### 3.2. R4F4-based peptides, including their cyclic and D analogues, exhibit strong antibacterial properties

We synthesised eight arginine-rich peptides to test their antimicrobial activity. They were tested against five bacterial strains, including the multidrug-resistant *P. aeruginosa* strain NCTC 13437. The *in vitro* findings contradict some of the *in silico* predictions, as certain potential antimicrobial peptides did not exhibit any effects on cultured bacteria. In the initial screening, R4F4 stood out as the most effective candidate, displaying lowest MIC and MBC values against both Gram-positive and Gram-negative bacteria (31.25→1000 µM). This is consistent with our previous report (ref.17). Due to its superior antimicrobial activity, we designed three chemically modified peptides based on the R4F4 backbone: a lipidated version with palmitoyl (R4F4-C16), a synthetic version with alternating L and D amino acids D-R4F4, and a cyclic analogue CP-R4F4 Lipidation reduced the antimicrobial activity of R4F4 (>1000 µM), while the incorporation of unnatural amino acids (D) and cyclisation enhanced its antimicrobial potential, resulting in MIC and MBC values (15.62→250 µM) which are closer to those of reference antibiotics **(Table 2)**. A lipopeptide comprising four arginine residues and a fatty acid with a 16-carbon chain was evaluated, demonstrating strong activity within a range of 62.5→1000 µM. This suggests that sequence reduction can be favourable in the context of lipidation.

**Table 2.**
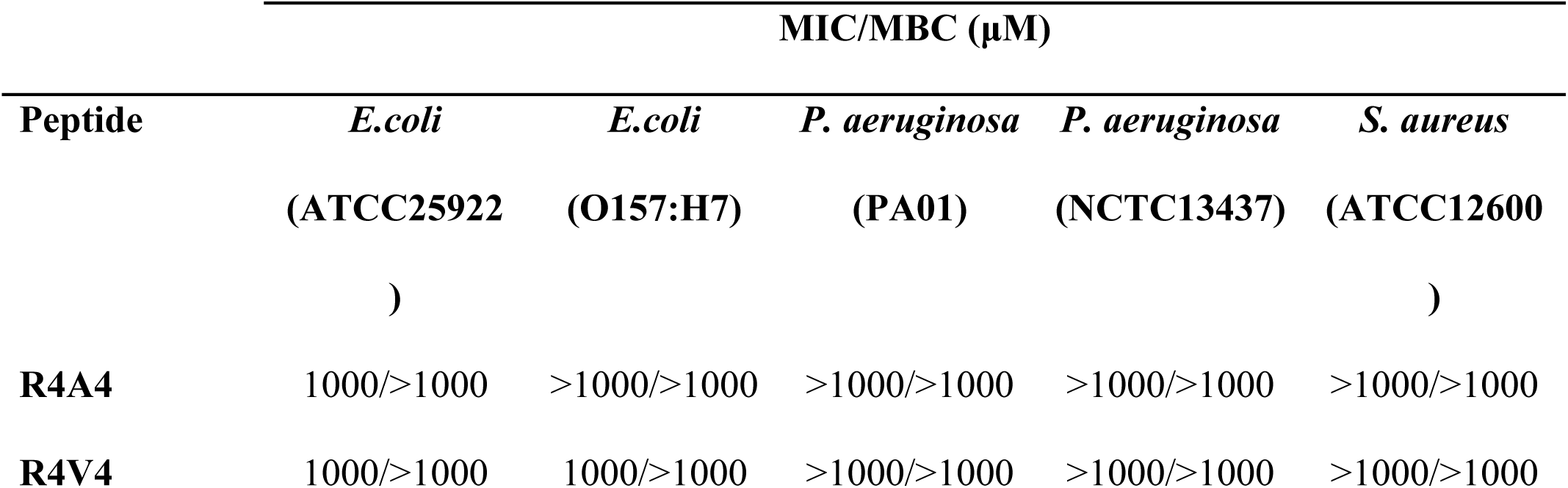
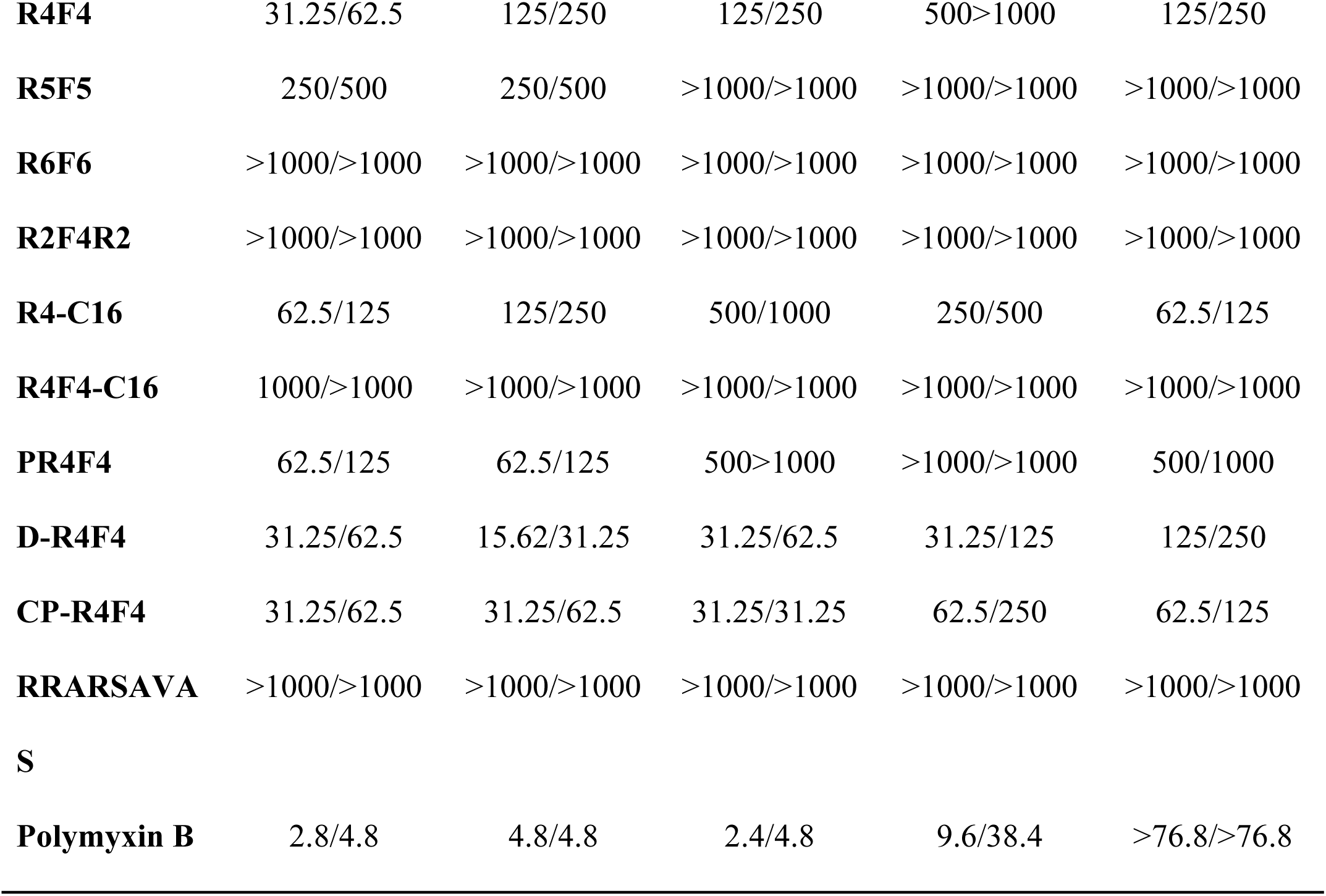
Antimicrobial activity of arginine-based peptides. The killing effects on bacteria were evaluated by determining the MIC and MBC values for both Gram-negative and Gram-positive bacteria exposed to arginine-rich peptides. A total of 5 x 10^6^ bacterial cells were incubated with a series of arginine-based peptides (0.03-1mM) in 96 well-plates, in triplicate, for 24 hours. The inhibition of bacterial growth was then assessed at 600 nm using an absorbance-based assay, with untreated controls as the baseline (100% growth).

### 3.3. Arginine-rich peptides have membrane-disrupting actions

To investigate the toxicity mechanisms of our arginine-rich peptides, we performed fluorescence-based experiments to measure changes in membrane permeability. Firstly, we employed a membrane potential sensitive fluorescent dye DiSC_3_(5) to evaluate the ability of peptides at 2× MIC or the highest evaluated concentration (1 mM) to depolarise the cytoplasmic membrane of *E. coli*. All peptides caused the migration of the dye to the extracellular environment, producing fluorescence signals. Greater effects were detected for *E. coli* treated with Triton X-100 and R4F4-C16 (**Figure 1A**). Then, fluorometrically, we evaluated if the peptides cause damage to the outer membrane using a lipophilic dye. The entry of NPN into the phospholipid layer resulted in substantial fluorescence in most treatments. The lipidated peptides showed higher uptake of the fluorescent dye (**Figure 1B**). Finally, we investigated the impact on the inner membrane of both *E. coli* and *S. aureus* strains using PI. The most active synthetic peptides showed higher incorporation of PI and consequently higher fluorescence levels, suggesting significant compromise of membrane integrity (**Figure 1C and D**).

**Figure 1.**
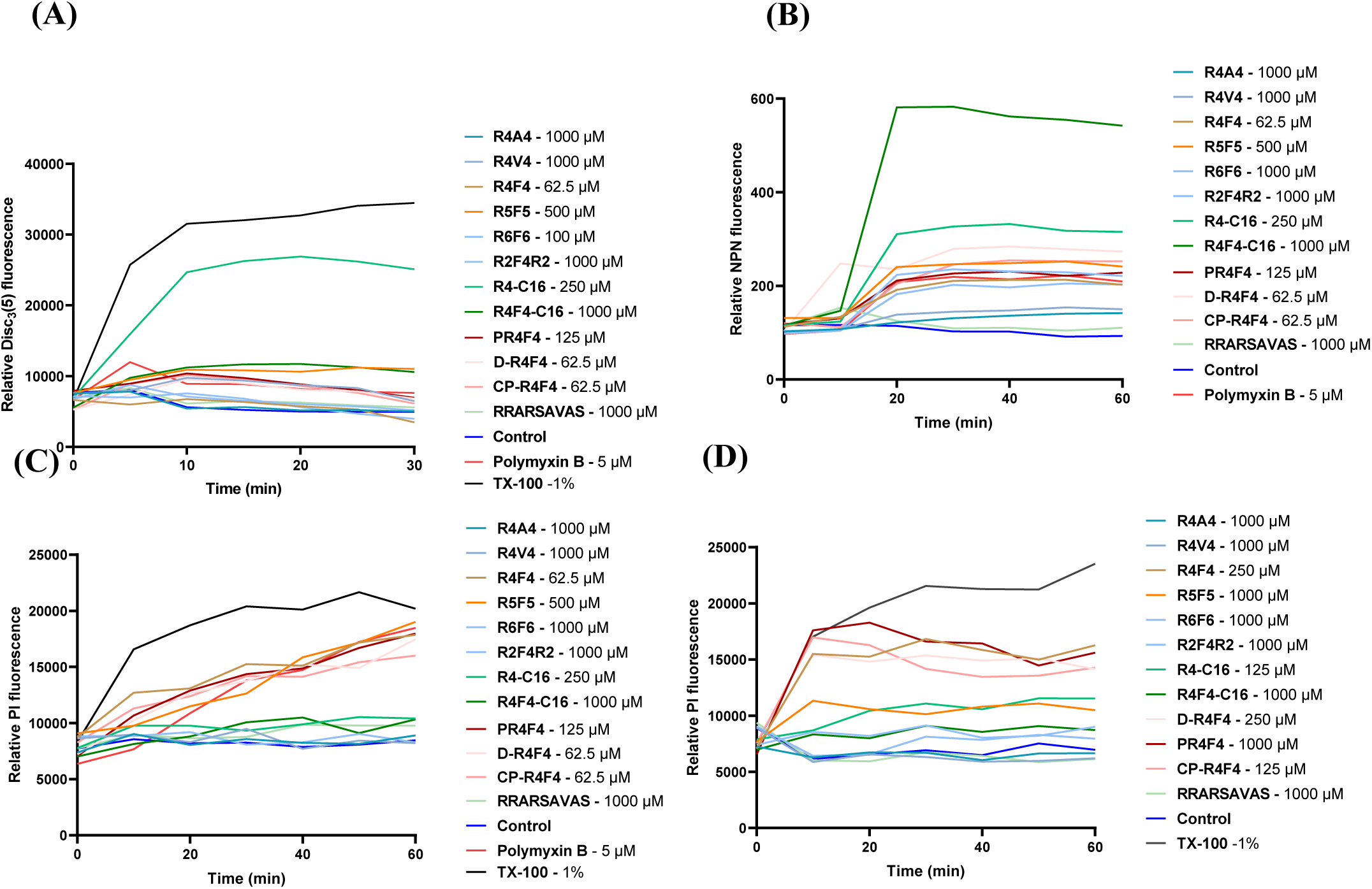
Membrane-targeting properties of arginine-rich peptides. **(A)** Cytoplasmic membrane depolarisation of *E. coli* ATCC 25922 was evaluated using 10 µM DiSC3(5). Increased fluorescence was detected for most of the peptide concentrations tested (2× MIC or 100 µM), except for R4C16, which exhibited significantly higher fluorescence levels compared to the positive control, TX-100. **(B)** Outer membrane permeabilisation profile of peptides against *E. coli* ATCC 25922. Mid-log phase cells were incubated with either 2× MIC or 1000 µM peptide for 1 h. Lipopeptides exhibited greater outer membrane permeabilisation than their analogue peptides. Inner membrane permeabilisation of **(C)** *E. coli* ATCC 25922 and **(D)** *S. aureus* ATCC 12600 after incubation with arginine-rich peptides. Propidium iodide uptake confirms the membrane-disruptive activity of R4F4, R5F5, R4-C16, PR4F4, D-R4F4 and CP-R4F4.

In addition to fluorescence microplate assays, we validated bacterial membrane damage induced by the most promising peptide candidates using microscopy-based imaging techniques. Initially, we monitored changes in the membrane permeability employing a two-colour nucleic acid staining fluorescence kit, which combines dyes that differentiate cells with either intact or compromised membranes. Cells with damaged membranes stain red, while intact membranes stain green. All cells treated in the negative control group (PBS) remained green throughout our experiments. Conversely, representative fluorescent images of bacteria incubated with Polymyxin B and the R4F4 peptide, as well as its modified analogues D-R4F4 and CP-R4F4 displayed mixed populations, with a high presence of red-stained cells indicative of damaged membranes (**Figure 2A**). To achieve three-dimensional and high-resolution visualisation of the membrane-disrupting action of the peptides at nanoscale, we used atomic force microscopy (AFM). The images revealed structural changes in the topography, roughness and integrity of bacterial membranes incubated with peptides. Clear leakage of intracellular content, indicating peptide-induced pore formation, was visualised (**Figure 2B**).

**Figure 2.**
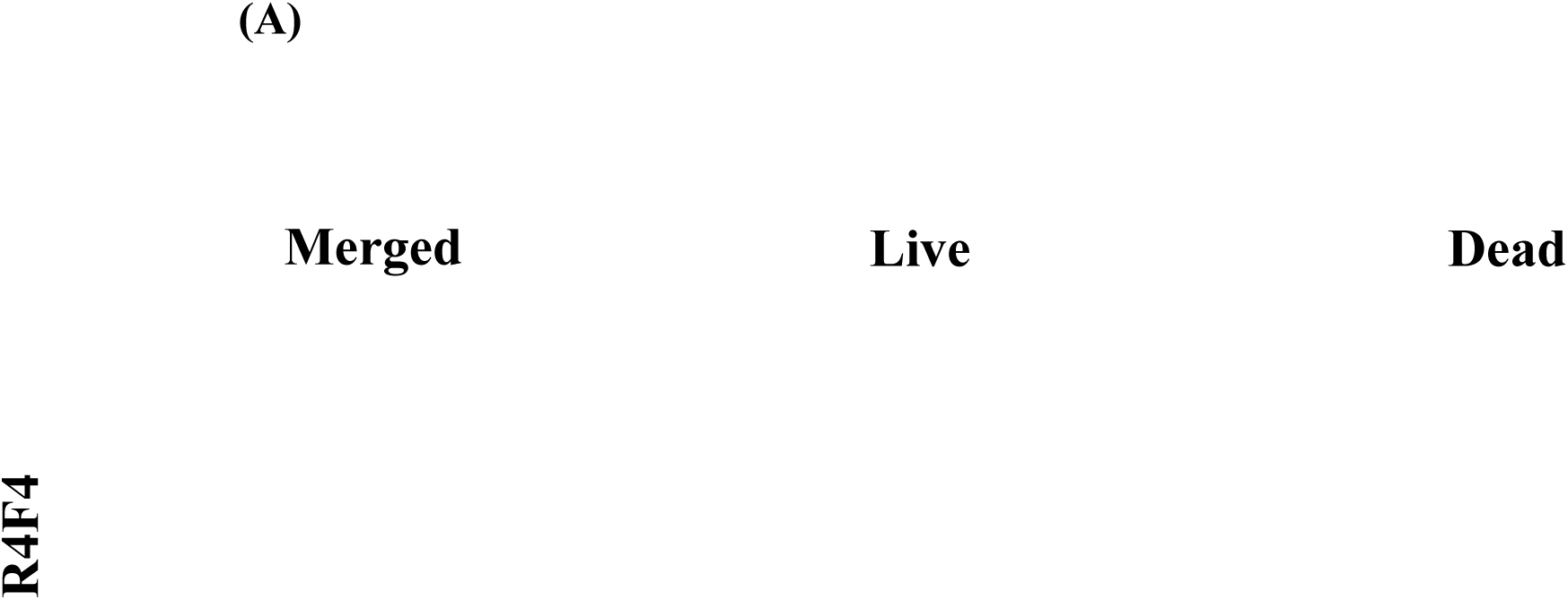
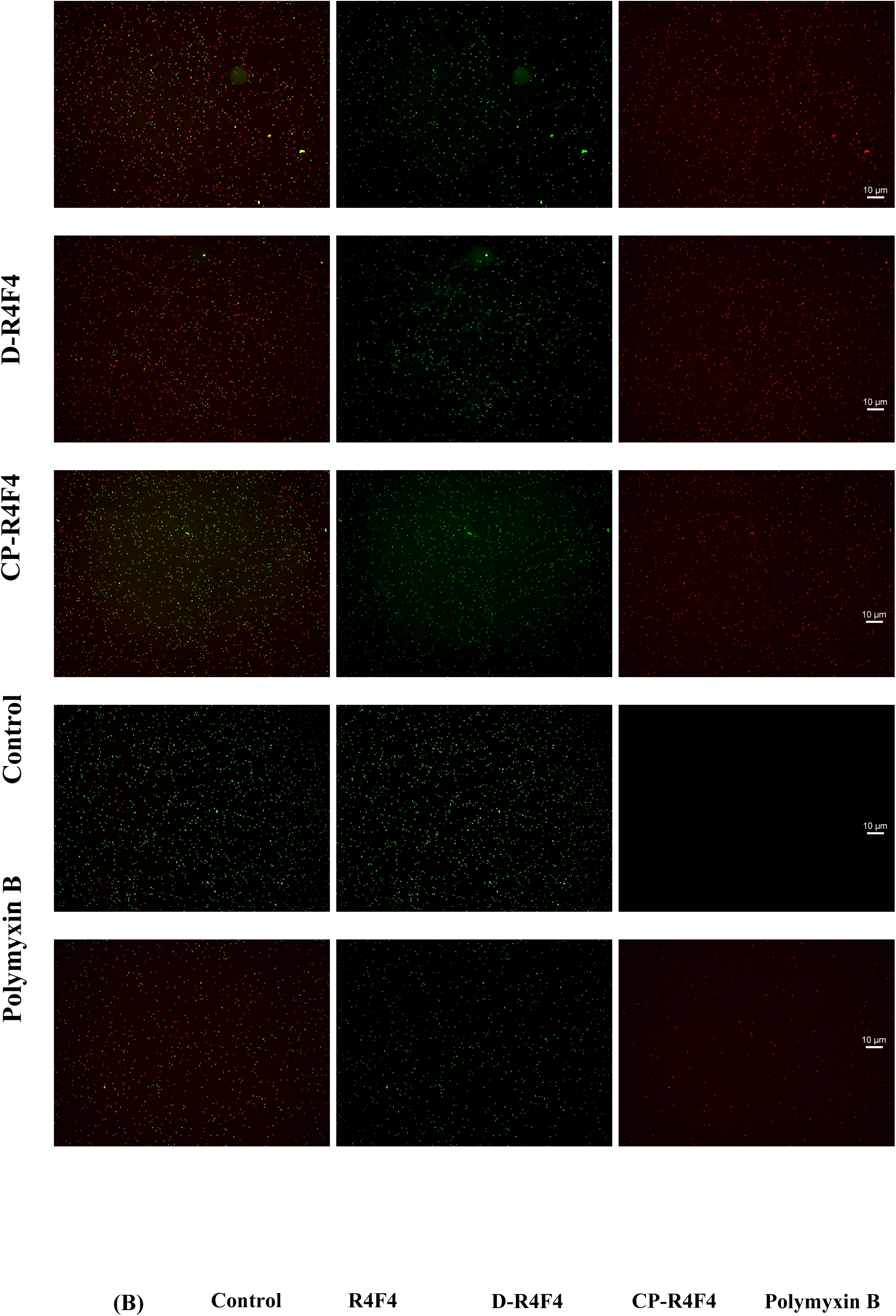
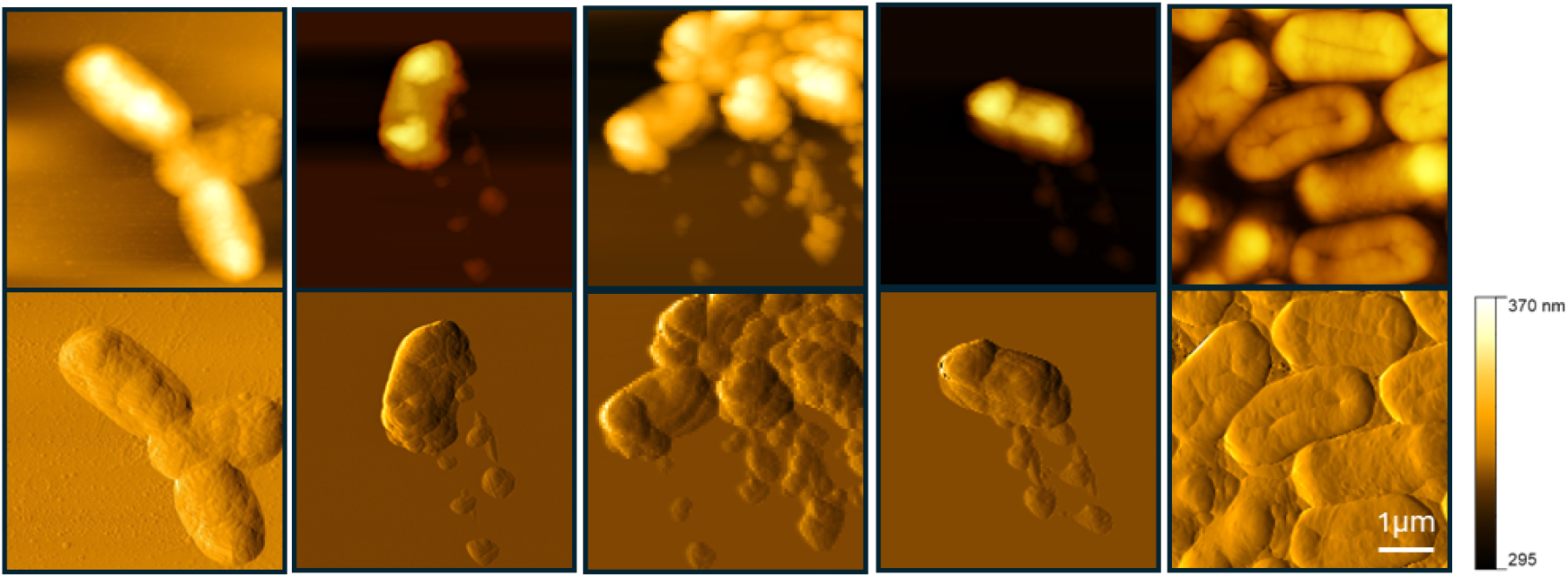
Arginine-rich peptides cause extensive damage to *E. coli* membranes. **(A)** Live/Dead cell panel showing the rapid effects of R4F4, D-R4F4, and CP-R4F4 on membrane integrity. Mid-log phase *E. coli* cells were incubated with 2× MIC, followed by staining with a SYTO 9/PI. Live and dead cells were stained green and red, respectively. Polymyxin B was added as a positive control. **(B)** Atomic force microscopy observations of *E.coli* confirmed the membranolytic effect of peptides, with leakage of cellular contents visible in the lower set of images. Experiments were based on three separate experiments.

### 3.4. D-R4F4 induces changes in the gene expression profile of *E. coli*

To further understand the molecular mechanisms underpinning the antimicrobial action of D-R4F4, we performed an RNAseq analysis of *E.coli*. D-R4F4 peptide was selected for its potent antibacterial activity. Mid-log phase cells were incubated with the peptide at MIC for 24 hours. Total mRNA was extracted from three independent biological replicates of bacteria treated with the peptide and compared to three control groups that were exposed to PBS. The heatmap generated through clustering analysis highlights the differential gene expression for both untreated and treated bacteria, while also revealing the consistent patterns among replicates **(Figure 3A)**. Among 9210 expressed genes, 1986 differentially expressed genes (DGEs) were significantly downregulated and 2038 upregulated genes in response to D-R4F4

**Figure 3.**
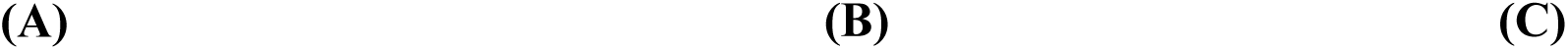
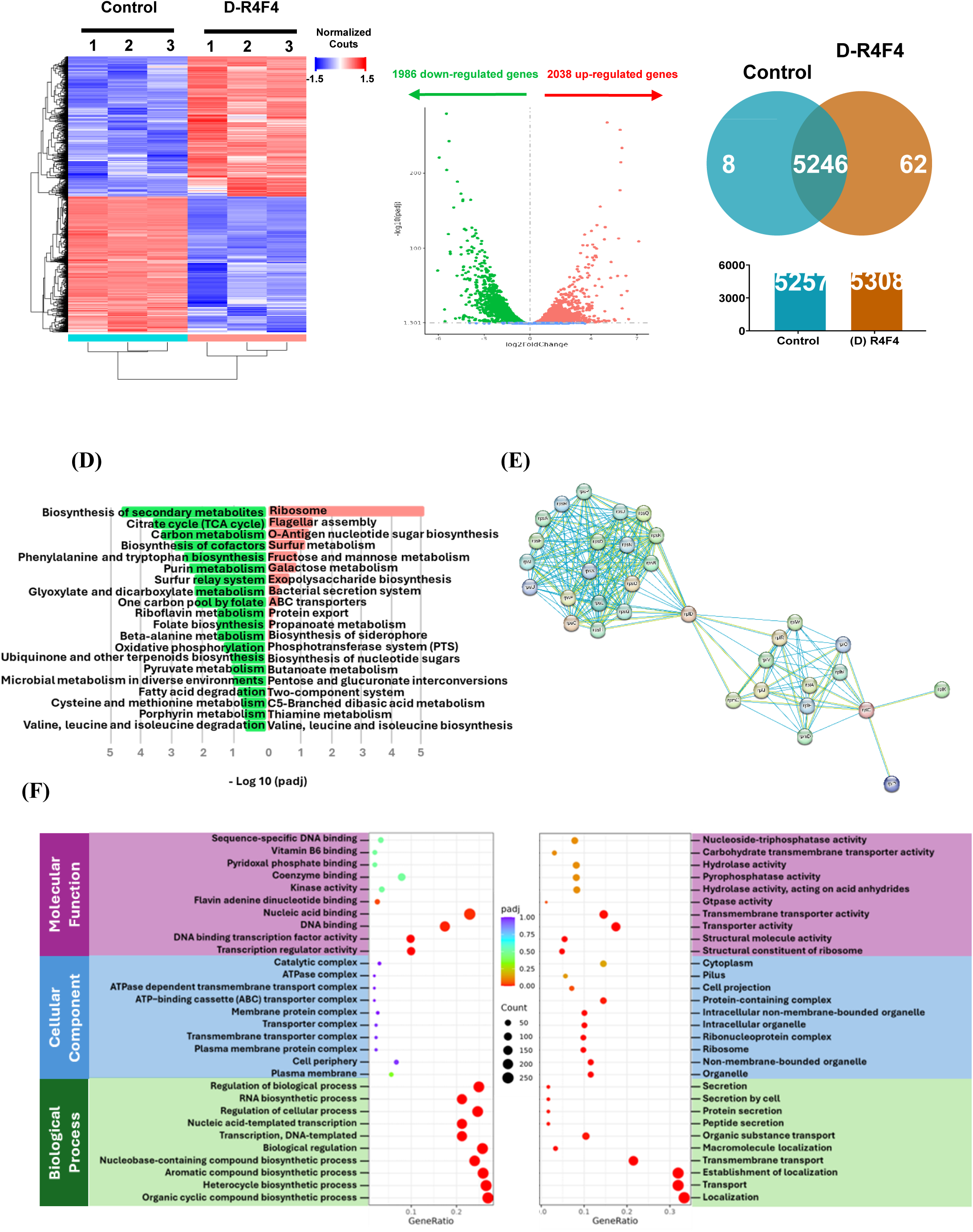
*E. coli* RNA sequencing data after D-R4F4 peptide exposure. **(A)** Hierarchical clustered heatmap depicting the relative expression of various genes compared with the negative control and D-R4F4 after 24h. **(B)** Volcano plot of the distribution of differentially expressed genes |log2(FoldChange)| > 1 and qvalue < 0.005. **(C)** Venn diagram of total mRNA expression levels in each group. **(D**) KEGG analysis showing downregulated (green) and upregulated (red) pathways. **(E)** Functionally associated network of upregulated ribosomal genes from STRING database. **(F)** GO enrichment of down- and up-regulated genes, shown in the left and right panel, respectively.

**(Figures 3B and 3C)**. KEGG (Kyoto Encyclopedia of Genes and Genomes) analysis revealed major alterations in gene expression, with downregulated genes primarily involved in the biosynthesis of secondary metabolites, the citrate cycle (TCA cycle), and carbon metabolism. In contrast, upregulated genes were associated with ribosome biogenesis, flagellar assembly, and O-antigen nucleotide sugar biosynthesis **(Figure 3D)**. Interestingly, genes associated with both the large (50S) and small (30S) ribosomal subunits, which formed highly interconnected expression clusters, were upregulated upon peptide exposure **(Figure 3E)**.

Gene Ontology (GO) enrichment analysis showed that downregulated genes were predominantly involved in RNA biosynthesis, regulation of biological and cellular processes **(Figure 3F)**. Among these, several key genes essential for *E. coli* survival under stress conditions were significantly downregulated. These included *rpoS*, a master regulator activated during environmental stress; *phoQ*, a membrane-associated sensor that modifies lipid A, thereby influencing outer membrane charge and susceptibility to antimicrobial peptides; *envZ* and *ompR*, which constitute an osmotic pressure sensing, two-component system that regulates porin expression (*OmpF/OmpC*), affecting membrane permeability; and *dnaA*, the initiator of chromosomal replication, whose reduced expression indicates potential impairment in DNA replication and cell division. In contrast, upregulated genes were mainly enriched in pathways related to transport, establishment of localisation, and localisation. These included *corA*, *pitA*, and *pitB*, which are involved in magnesium and phosphate ion transport; *ybtP*, *ybtQ*, and *ybtX*, which encode components of the siderophore-dependent iron uptake system; and *fruA*, *srlA*, *srlB*, *malF*, *malG*, and *malM*, which are responsible for the transport of sugars such as fructose, sorbitol, and maltose.

### 3.5. Antibacterial arginine-rich peptides enhance the generation of reactive oxygen species (ROS)

Although the membrane is the primary target for cationic peptides, other effects can also contribute to their killing efficacy and potency. To expand our understanding of the mechanisms by which arginine-rich peptides induce the death of clinically relevant bacteria, we monitored changes in intracellular ROS content using a sensitive cell-permeable dye. These alterations were detected through both fluorescent reads and microscopic imaging. Peptides with higher antimicrobial properties showed an increased ROS generation in a concentration-based response manner (**Figure 4A**). Among all treatments, R4-C16, D-R4F4 and CP-R4F4 showed the highest modulation of ROS levels in *E. coli and S. aureus*. A representative image of peptide-treated *E. coli* is shown in **Figure 4B**. Additionally, exposure of *E. coli* to D-R4F4 resulted in the downregulation of a set of genes related to oxidoreductase activity **(Figure 4C),** with over 80 differentially expressed genes (DEGs) being downregulated in total. **Figure 4D** highlights downregulated DEGs involved in oxidative stress (showing only proteins with interactions).

**Figure 4.**
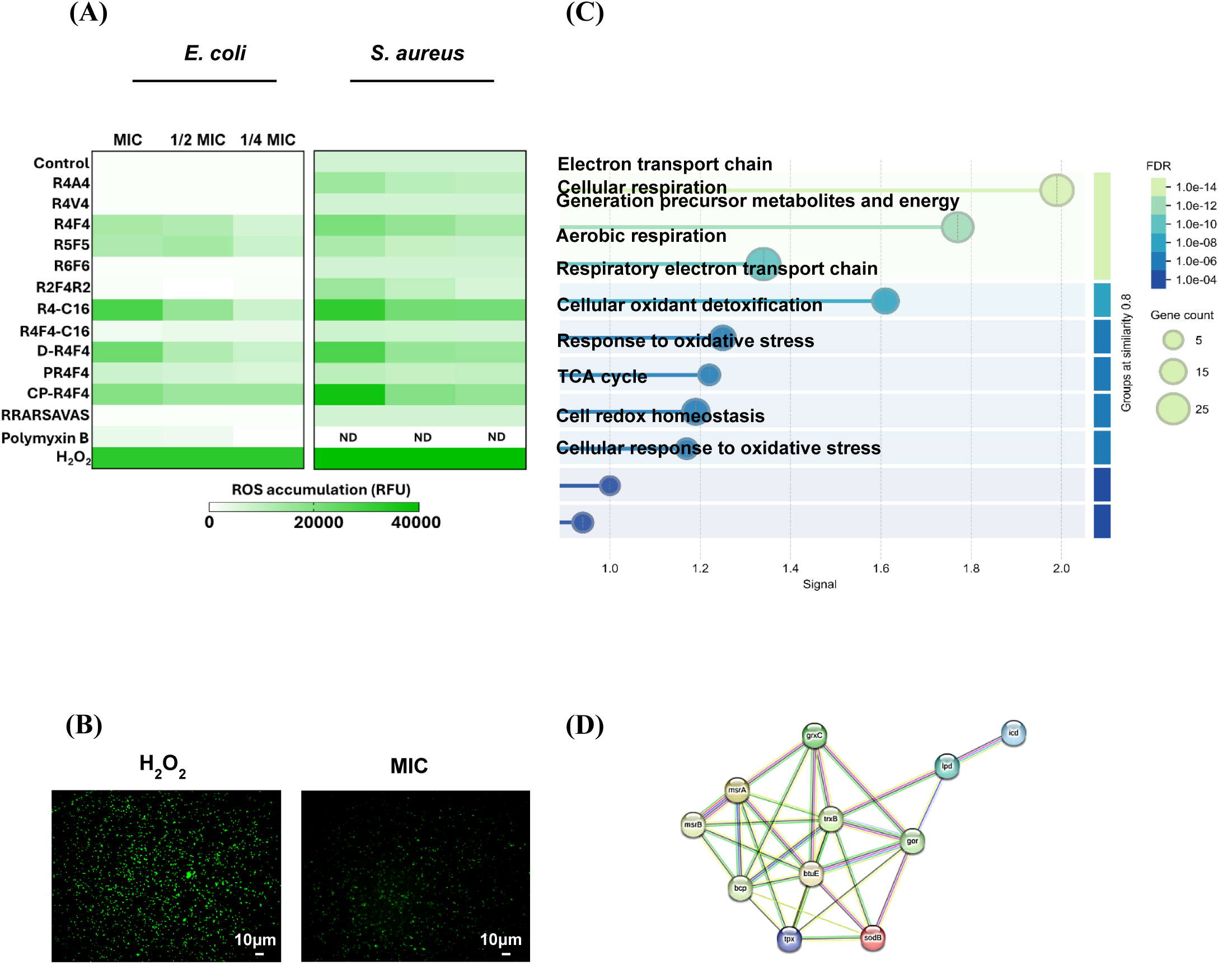
**Arginine-rich peptides increase the total reactive oxygen species (ROS)**. **(A)** Oxidative stress levels in *E. coli* and *S. aureus* after exposure to peptides at MIC for 4 h at 37°C. **(B)** Fluorescence images show the impact of D-R4F4 on *E. coli* intracellular ROS accumulation. **(C)** GO terms for molecular function associated with downregulated differentially expressed genes (DEGs) involved in oxidoreductase activity in *E. coli* after D-R4F4 peptide treatment for 24 h. **(D)** Protein-protein interaction network for downregulated DEGs identified in response to oxidative stress (STRING database).

### 3.6. Arginine-rich peptides inhibit biofilm formation and promote its disruption

Early-stage biofilm formation in the presence of the most active arginine-rich peptides was evaluated using the crystal violet (CV) microtiter plate assay. All three peptides showed anti-biofilm properties, with a strong capacity to limit biofilm development. Overall, the biofilm biomass stained by CV was significantly reduced in the presence of peptides. D-R4F4 and CP-R4F4 exhibited the lowest levels of biofilm formation at the lower concentrations (**Figure 5A**).

**Figure 5.**
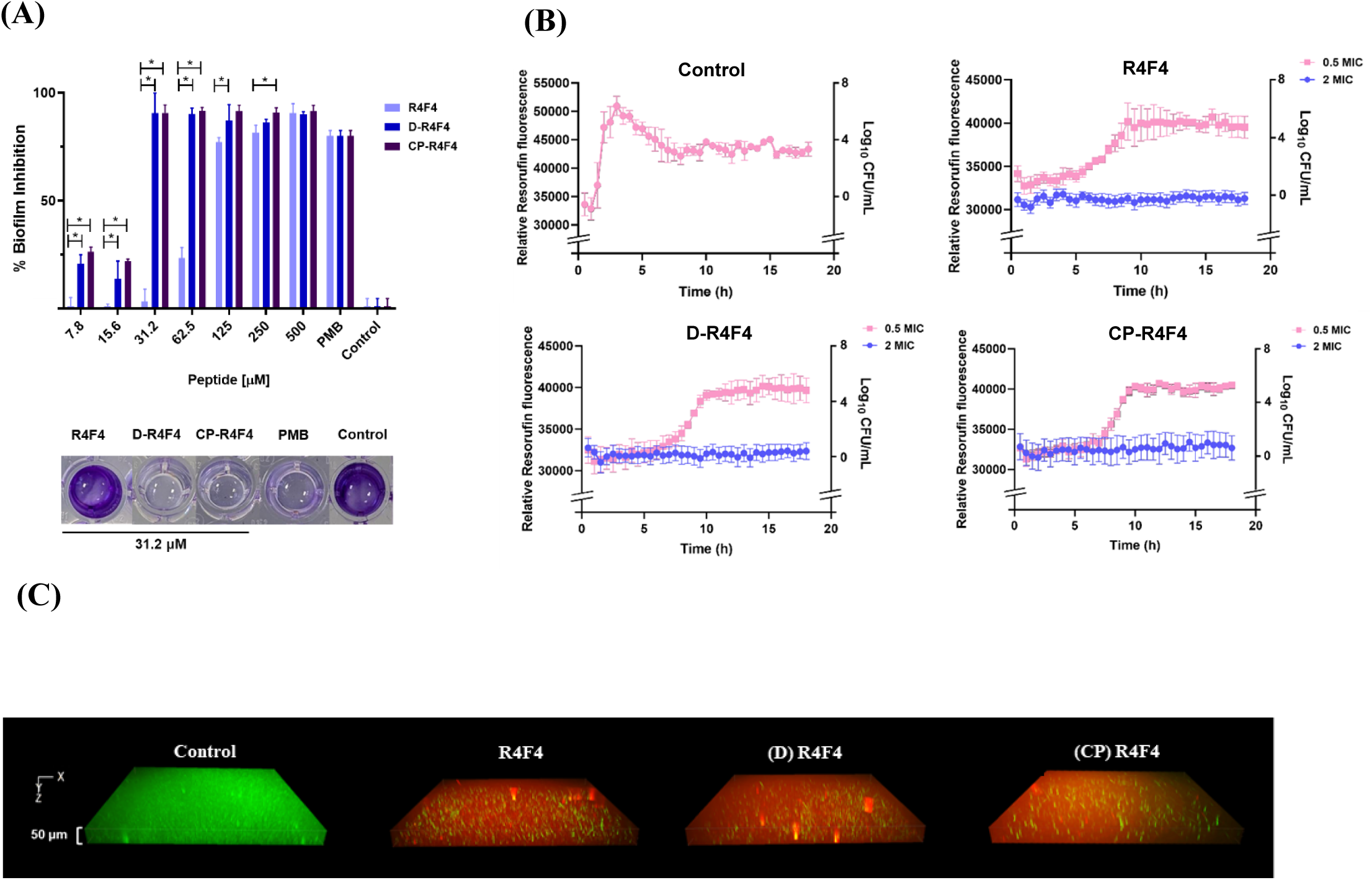
Antibiofilm activity of R4F4, D-R4F4, and CP-R4F4. Peptides were evaluated in both stages of biofilm formation: **(A)** Inhibition and **(B-C)** Disruption. **(A)** *P. aeruginosa* was incubated with peptides at concentrations ranging from 0 to 500 µM for 24 h. Biofilm mass was stained with 0.1% crystal violet (CV) and measured at OD_597_. Data represent the percentage of treatment relative to the growth control (mean ± standard deviation), which were calculated from three independent experiments: (*) p<0.001, by one-way analysis of variance (ANOVA) with Tukey’s post hoc tests. PMB refers to Polymyxin B at its MIC value. **(B)** Kinetic plot showing the metabolic activity of the biofilm when treated with 0.5 or 2× MIC of peptides. Mature biofilms were allowed to grow in M63 supplemented for 48h. Peptides were added in fresh MH broth containing 100 μg/mL resazurin. Fluorescence was monitored at λ_ex_= 520 nm / λ_em_= 590 nm over 18h. **(C)** Microscopic imaging of 3D biofilm. Bacterial viability staining with SYTO9/PI confirmed biofilm damage upon peptide exposure, with dead and viable bacteria appearing in red and green, respectively, after 3h incubation.

The effect of peptides on pre-established biofilms was also assessed using a resazurin-based viability assay. In the control group, non-fluorescent resazurin was reduced to resorufin by metabolically active bacteria, resulting in high levels of fluorescence (**Figure 5B**). Similarly, at 0.5 MIC, all three peptides showed a profile with a high presence of viable cells. However, the biofilms were significantly disrupted when incubated with 2×MIC of R4F4, D-R4F4, and CP-R4F4. The anti-biofilm properties of arginine-rich peptides were confirmed through confocal microscopy using membrane integrity-sensitive DNA-binding dyes. A dense green bacterial community was observed in the control group (**Figure 5C**), while populations exhibiting both green and notably red signals were observed in the peptide-treated groups, confirming the potential of R4F4 and its analogues to disrupt established biofilms.

### 3.7. Chemically modified R4F4 analogues retained activity in protease-rich environments

Peptides rich in positively charged amino acids are prone to structural degradation in proteolytic environments such as serum. Therefore, we assessed the antimicrobial activity of arginine-based peptides incubated with both trypsin and serum. Changes in the ability of peptides containing natural amino acids to kill both Gram-positive and Gram-negative bacteria were observed, particularly for R4F4. The proteolytic action of trypsin alters the activity of R4F4 but does not affect the micromolar efficacy of R4F4-C16, D-R4F4, and CP-R4F4 (**Table 3**). Polymyxin B also showed stability in the presence of serum and trypsin.

**Table 3.**
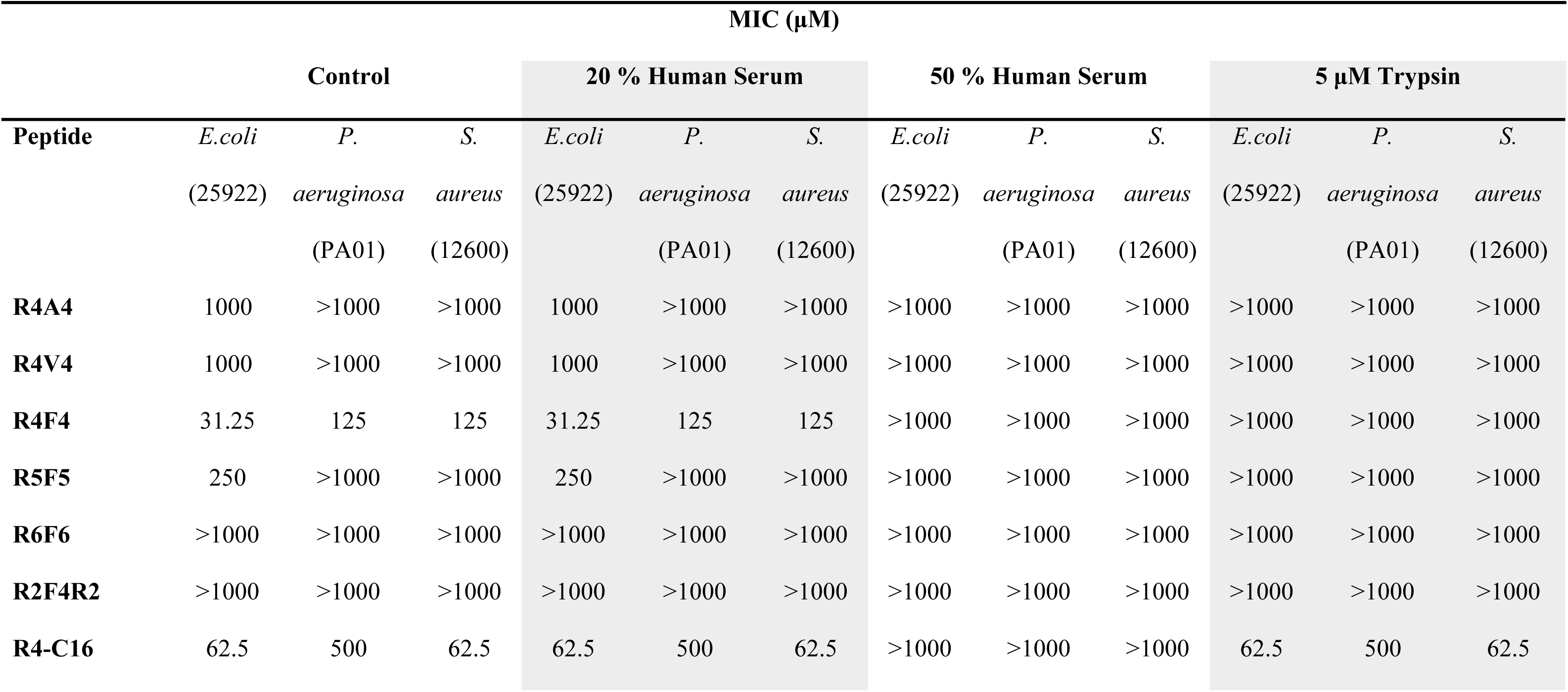
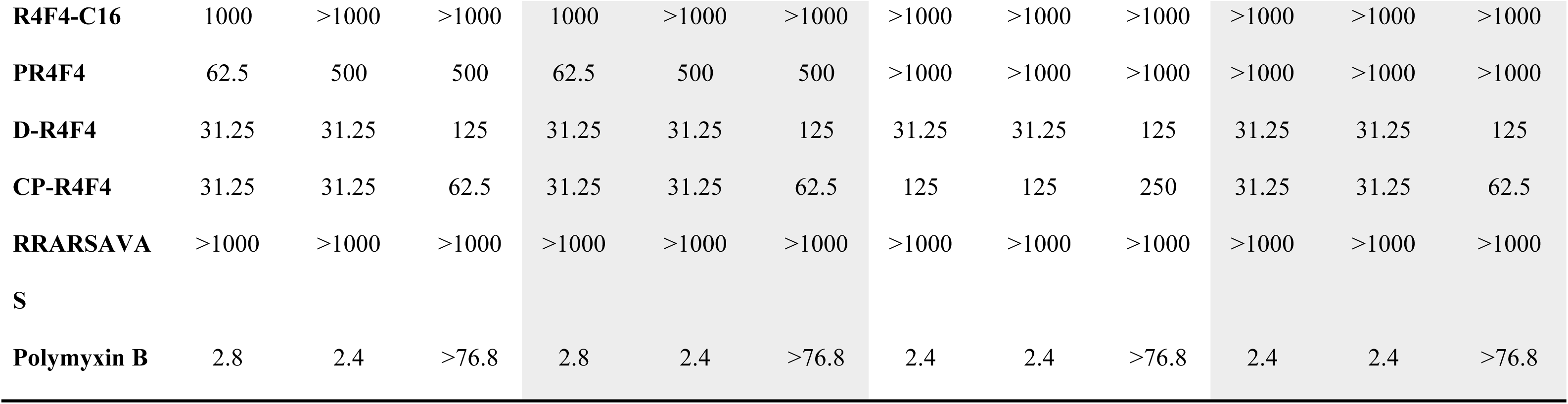
Antimicrobial activity of arginine-rich peptides in presence of serum and trypsin. Different concentrations of arginine-based peptides were incubated with serum and trypsin and the MIC was determined.

To better understand the alterations in the MIC values of R4F4 and confirm the stability of chemically modified analogues D-R4F4 and CP-R4F4, we conducted a mass-spectrometry-based (MS) analysis of these peptides in the presence of serum and purified trypsin. The R4F4 peptide was degraded into smaller fragments when incubated with trypsin. The peak with a retention time of 5.89 minutes, corresponding to the characteristic molecular weight of R4F4, disappeared under proteolytic conditions (**Figure 6**). In contrast, the chromatographic profiles and corresponding spectra of (D-R4F4 and CP-R4F4) demonstrated the stability of these molecules, which appeared at the same retention time in both the absence and presence of proteases. The molecular weight of these modified peptides corroborated their chemical resistance to the enzymatic action of trypsin and serum proteases.

**Figure 6.**
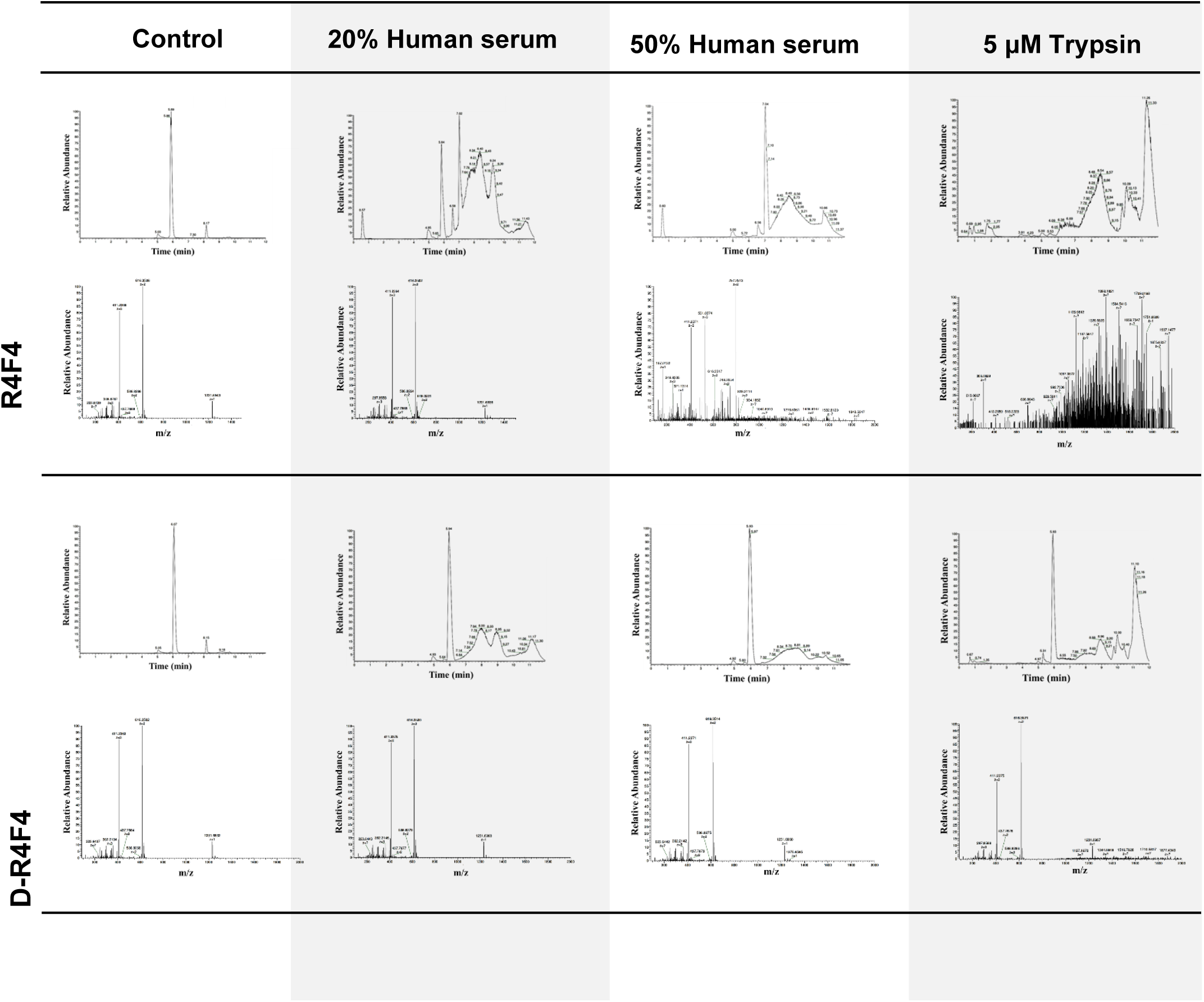
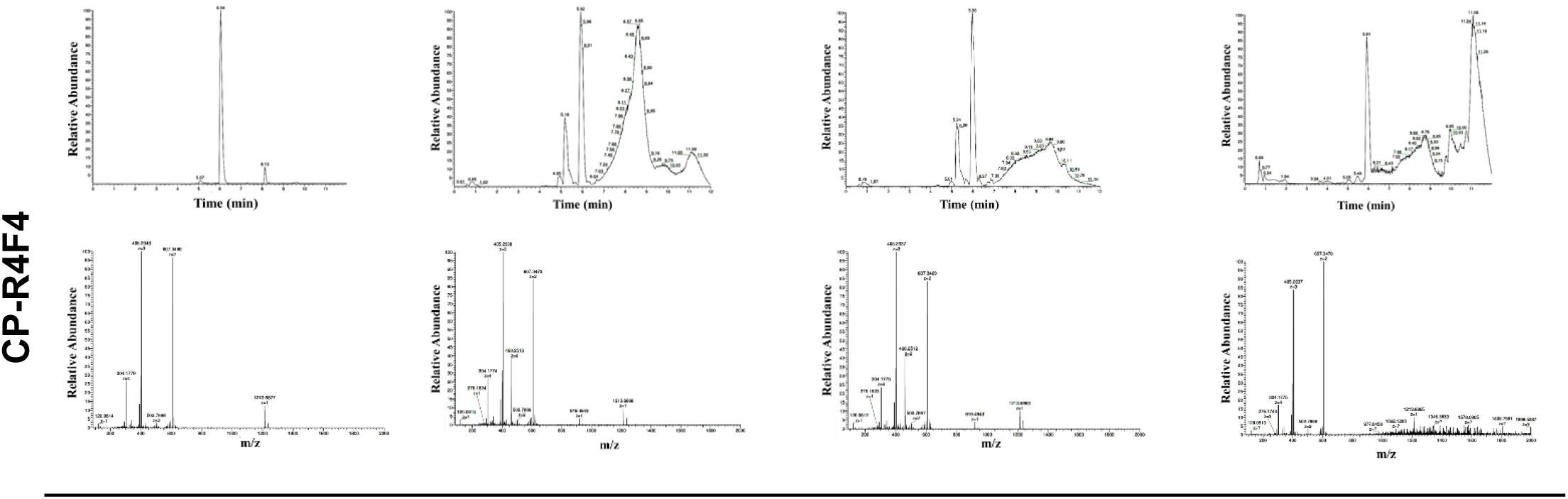
Chemically modified R4F4 analogues remain stable under proteolytic conditions. Representative LC chromatograms (upper graphs) and mass spectra (bottom graphs) of arginine-rich peptides. Synthetic molecules were incubated with serum and trypsin and analysed by LC-MS.

### 3.7. Snapshot of peptide-induced toxicity

In the final assessment, we evaluated the cytocompatibility of arginine-rich peptides using two cell models: anucleate cells (hRBCs) and nucleated cells (fibroblast L929). Peptides lacking antimicrobial activity or exhibiting low predicted potential also showed minimal toxicity toward both cell types. However, lipidated peptides exhibited the highest toxicity, indicated by the greatest release of haemoglobin from hRBCs and the lowest fibroblast viability, as determined by the MTS assay. R4F4, D-R4F4, and CP-R4F4 all demonstrated toxicity at concentrations higher than those required for antimicrobial effects, suggesting lower chances of adverse effects (**Figures 7A-B**).

**Figure 7.**
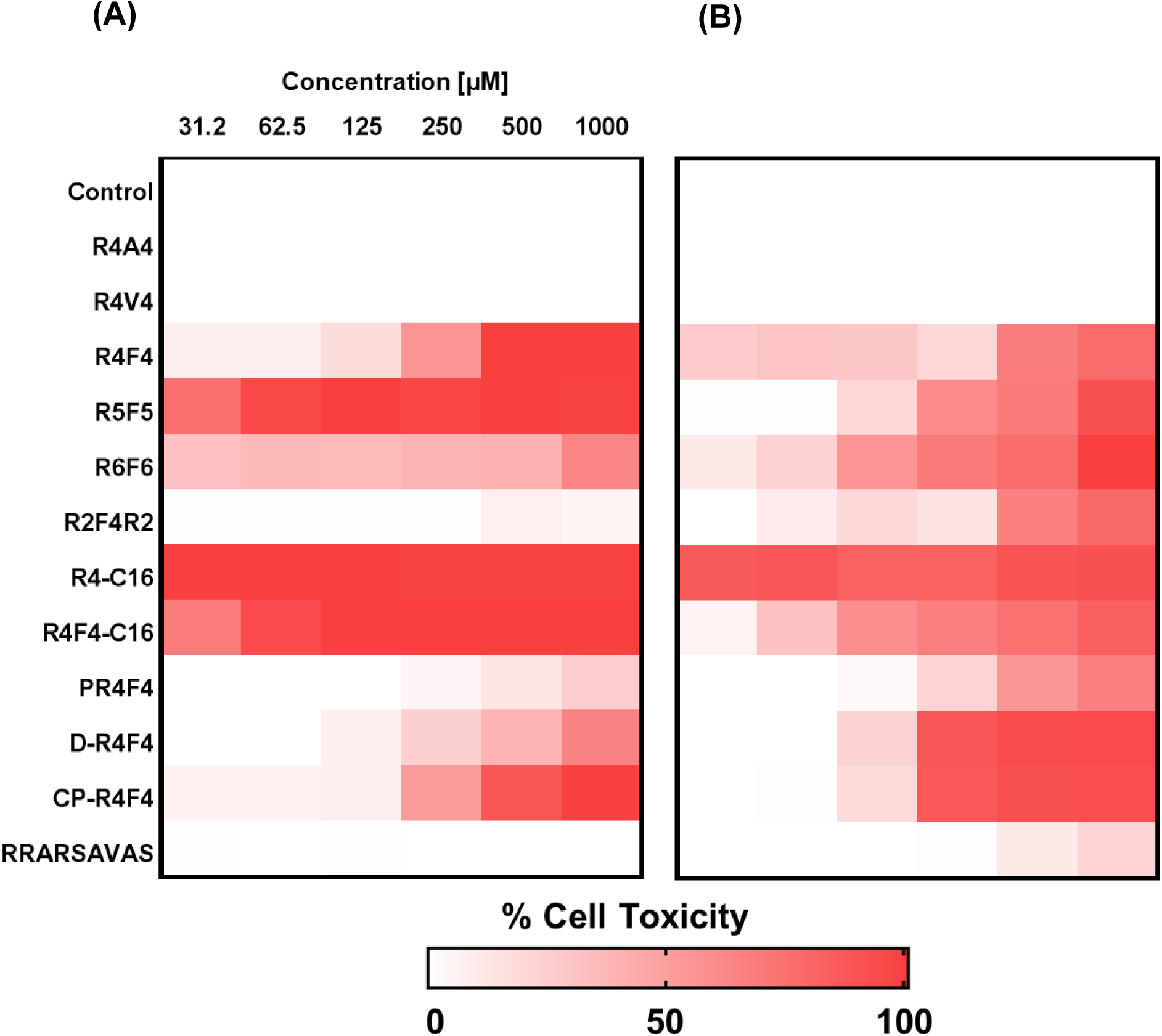
Haemolytic and cytotoxicity profiles of arginine-rich peptides. **(A)** Human RBC and **(B)** L929 fibroblasts cell lines were used to screen the *in vitro* toxicity of peptides. Lipidated peptides showed the highest toxicity toward both human cell types. The heatmap was generated from an average of three independent experiments.

## 4. DISCUSSION

The effectiveness of clinically relevant agents in treating bacterial infections is undermined by rising levels of antimicrobial resistance, largely driven by both the mis- and overuse of existing antibiotics [27]. Currently, antimicrobial peptides form part of the arsenal in treatment regimens for multidrug-resistant bacteria [28,29]. However, there remains a pressing need to further develop and design novel antimicrobial peptides which are clinically applicable and effective against a range of pathogenic bacteria [30]. Peptide-based antibiotics are usually rich in positively charged amino acids, which interact strongly with both the negative and hydrophobic components of bacterial membranes [31,32]. Chemically, peptides are highly versatile, providing ample opportunities for modification and optimisation, which enhances their potential for effective clinical translation [33,34]. Exploring the combinatorial library of peptides can broaden the range of options for combating bacterial infections.

For many years, the discovery of therapeutic antibiotic peptides has been conducted through the systematic screening of molecules derived from natural sources (e.g. plants, animals), often requiring rigorous and time-consuming steps at the laboratory bench [35]. More recently, the development of AI (artificial intelligence)-driven prediction tools has accelerated the screening process, enabling the identification of potential candidates in a shorter time frame with higher chances of success [36,37]. In this study, we used *in silico* tools to predict the antimicrobial and haemolytic properties of eight arginine-rich peptides composed of natural amino acids. Discrepancies were observed between the predictions made by AMPFun and Antimicrobial Peptide Scanner v .2. These discrepancies have also been reported in previous studies [38,39], highlighting the importance of reliable peptide sequence datasets and corresponding in vitro results for developing more robust predictive models, especially for short de novo designed peptide sequences, where databases of natural (longer, less regular) will not provide an accurate description. Additionally, this underscores the necessity of experimental assays to validate and, in essence, ground-truth *in silico* results. To confirm this initial computational assessment, the peptides were synthesised, purified to >95% purity, and tested against a set of Gram-positive and Gram-negative bacterial strains.

Our *in vitro* data were more consistent with the results obtained from Antimicrobial Peptide Scanner v .2. However, some peptides predicted to exhibit antimicrobial activity did not exhibit any activity even at the highest tested concentration. Although *in silico* strategies can assist in identifying potential candidates and conserving resources by focusing *in vitro* assessments on those with higher probabilities of effectiveness, many classification-based methods can still fall short in identifying potent structures, as we have demonstrated in our study. For example, although R5F5 and R6F6 were predicted to have antimicrobial activity by two web servers, they did not exhibit any *in vitro* activity against any of the strains assessed. Adding additional data to freely available peptide databases, along with the use and validation of available in silico tools, is essential in enhancing scoring functions, refining existing models, and developing more accurate methods, as suggested in the literature [40,41].

The analysis of the antimicrobial properties of linear peptides containing natural peptides showed that arginine combined with phenylalanine has the strongest *in vitro* effect. That phenylalanine can contribute to the antibacterial activity of cationic peptides is supported by other studies [38,42] However, when challenged with trypsin ( a protease capable of degrading peptides), R4F4 was fragmented and lost its potency, which can lessen its clinical applicability. Our results have been corroborated by other studies, which have shown a reduction in the activity of cationic peptides in the presence of proteolytic enzymes [43,44]. Thus, these findings highlight the need to optimise the biochemical structure of this template to make it more attractive for future drug development.

Due to challenges in tested protease-rich environments, we chemically modified the peptide R4F4 to assess stability. Attaching lipid tails to peptides to address these stability concerns is a widely applied strategy in various antimicrobial peptide templates [45,46]. Although this modification conferred good stability, the analogue (R4F4-C16) displayed off-target activity, showing a preference for hRBCs over bacteria. This example emphasises that the addition of a lipid tail may not equally benefit all peptide sequences and that careful examination is necessary before implementation. In agreement with this, other investigations have highlighted the complexity of fine-tuning toxicity and activity. Fatty conjugation can result in higher levels of haemolysis or toxicity and alter self-aggregation behaviour [45,47]. In this study, we designed two additional chemically modified versions of R4F4: a cyclic version (CP-R4F4) and an analogue with alternating L and D amino acids (D-R4F4). Both strategies ensure good stability and cytocompatibility, with antibacterial activity in the micromolar range. Positive outcomes have been reported by other authors [48,49]. These strategies were also adopted before leading to the development of commercially available antibiotics [50,51].

The effects of cationic peptides have traditionally been explained through their interaction with membranes, leading to pore formation and leakage of cellular components [52,53]. Our analysis showed that our peptide candidates possess membrane-destabilising properties, compromising the integrity and functionality of the bacterial membrane. However, this is not the sole mechanism of action, as our complementary investigations revealed the involvement of multiple additional processes. The fluorescent analysis of peptide-treated bacteria revealed a significant increase in ROS levels, which can cause damage and lead to bacterial death. Elevations in the generation of ROS have been reported for other AMPs [54,55]. In addition, D-R4F4 induces intracellular changes at the molecular level, as evidenced by our global transcriptional analysis of the response in *E. coli* incubated at peptide MIC. Interestingly, exposure to the synthetic peptide resulted in downregulation of several oxidoreductase-related genes, including *msrA*, *msrB*, *sodB*, and *grxC/grxD*, which indicates possible disruption of cellular detoxification pathways. These combined findings expand the understanding of the modulatory effects of arginine-rich peptides, which can act through multiple mechanisms, thereby potentially minimising the chance of bacterial resistance development.

Although arginine-rich peptides are well-established as templates for designing antibiotics and effective carriers owing to their cell-penetrating properties [56], the dynamic gene expression response in *E. coli* after peptide exposure has not been thoroughly investigated. A few studies via metabolomics [57,58], proteomics [59] and transcriptomics [60–63] have looked at this angle, although it is underexplored. Our analysis captured distinct gene expression profiles in both control and peptide-incubated cells. The mechanism underlying these transcriptional alterations remains unknown, but it is likely linked to membrane perturbations and increased permeability induced by the peptide, as intracellular signalling or specific binding to intracellular targets cannot be ruled out. To deepen our understanding of these molecular changes, tracking the D-R4F4 peptide and incorporating a biophysical perspective could provide a clearer picture of how the peptide is changing the transcriptomics landscape.

The peptide-induced transcriptional response was marked by the downregulation of pathways involved in the biosynthesis of secondary metabolites, the Krebs cycle, and carbon metabolism. The disruption of energy metabolism has similarly been observed in other RNA- and metabolite-based studies [58,61]. The changes also encompassed transcriptional activity and motility, as evidenced by higher transcript levels of ribosome-associated genes and elevated expression of flagellar assembly components. In line with this, a previous study reporting the incubation of *E. coli* with a cationic frog-derived peptide, known as magainin I, demonstrated a substantial increase in ribosome biogenesis and translation activity [64]. Bacterial motility can be modulated to facilitate exploration of more favorable environments [65]. The changes observed in our study indicate a substantial investment in intensified flagellar activity, possibly reflecting an adaptive response to evade the presence of D-R4F4, a phenomenon frequently observed when bacteria experience stress conditions [66].

Gene expression was down-regulated in the following functional areas: RNA biosynthesis, regulation of biological and cellular processes. Several critical stress-responsive and regulatory genes in *E. coli* incubated with D-R4F4 showed significantly lower levels of transcripts compared to untreated cells. In this context, we observed reduced expression of *rpoS*, which has implications for the susceptibility of *E. coli* to peptide-induced stress. On the other hand, increased membrane permeability and susceptibility are aligned with the suppression of *phoQ* and the downregulation of envZ and *ompR*. These regulations disrupt lipid A modifications in the outer membrane, compromising charge adjustments and impairing the expression of porins involved in membrane permeability, respectively. The reduced expression of *dnaA* indicated disruption in the replicative functions, hindering the survival under peptide-induced alterations. Overall, these transcriptional changes evidence the multifaceted impact of arginine-based peptides on bacterial survival mechanisms.

The *E. coli* response to stress induced by D-R4F4 was also characterised by transcriptional adjustments in genes linked to transport and localisation processes. As demonstrated in our study, this peptide destabilises bacterial membranes, leading to ion leakage similar to the membranolytic action of other cationic AMPs [67]. The upregulation of magnesium and phosphate ion transport (*corA, pitA, pitB*) is potentially a molecular strategy to restore ionic balance and homeostasis. We also found the upregulation of siderophore-dependent iron acquisition genes (*ybtP, ybtQ, ybtX*) are upregulated, possibly as part of a stress response to counteract oxidative damage caused by D-R4F4. *E. coli* survival relies heavily on iron, which is essential for facilitating enzymatic reactions and supporting crucial metabolic pathways [68]. Transcriptional alterations in order to increase iron uptake were also previously observed in *E. coli* responses to scorpion venom-derived peptides [69]. In addition, the elevated expression of sugar transport systems (*fruA, srlA, srlB, malF, malG, malM*) suggests an adaptive metabolic shift aimed at sustaining energy production. This aligns with impairments in the functioning of the TCA cycle, forcing *E. coli* to prioritise the uptake and use of readily available sugars to support to sustain basic and essential metabolic functions. In summary, these molecular changes illustrated the bacteria’s attempt to survive in the presence of this membranolytic short peptide by mobilising transport systems and optimising resource acquisition. However, such strategies might be insufficient to overcome the multifactorial stress and multimodal mechanisms exerted by this peptide, particularly in higher concentrations, which include membrane disruption and oxidative damage.

R4F4, and especially its two more potent analogues, CP-R4F4 and D-R4F4, can be promising alternatives for tackling biofilm-related infections. These peptides demonstrated dual action in both early-stage and mature biofilms. As evidenced by our results, they can inhibit biofilm formation and also adversely affect cells within existing biofilms. The transcription of genes associated with biofilm activity was also modulated by D-R4F4. Some upregulated genes, such as *pdeD* and *flhC/flhD*, are known to suppress biofilm formation. Conversely, the downregulation of *barA*, *uvrY*, and *rpoS* is also associated with impaired regulatory pathways essential for biofilm development. Earlier studies have demonstrated that antibiofilm peptides can synergise with conventional antibiotics, enhancing their antimicrobial properties [70,71]. Thus, a future avenue of this work could be to explore the interaction of arginine-rich peptides and antibiotics, which could be beneficial in the context of chronic infections, where biofilms pose significant challenges to existing treatments [72].

## 5. CONCLUSIONS

In conclusion, we computationally and experimentally mined cationic arginine-rich peptides for the selective activity on bacteria without inducing significant toxicity to two human cell lines. AI tools demonstrate significant potential in identifying novel candidates; however, shortcomings are also found in this approach. The combination of arginine and phenylalanine enhances the interaction with bacterial membranes and improves antimicrobial impacts. Incorporating D-amino acids or cyclisation aids in preventing proteolytic degradation, although lipidation did not result in better activity or reduced toxicity. Overall, this study provides insights for the future engineering of cationic, arginine-rich peptide sequences for drug development and their efficacy in more realistic environments, such as the presence of proteases. The transcriptomic perspective adds molecular level insight into the action of cationic peptides, moving beyond the simplistic view of their membrane-disruption mechanism. This approach can facilitate the identification of potential targets and pathways for developing future combinatorial antibiotic therapies.

## Declarations

Funding: IWH was supported by EPSRC Fellowship grant (reference EP/V053396/1)

Competing Interests: No competing interests Ethical Approval: None required

Sequence information: None to report

## REFERENCES

1. Ahmed, S.K.; Hussein, S.; Qurbani, K.; Ibrahim, R.H.; Fareeq, A.; Mahmood, K.A.; Mohamed, M.G. Antimicrobial resistance: Impacts, challenges, and future prospects. *Journal of Medicine*, Surgery, and Public Health 2024, 2, 100081, 10.1016/j.glmedi.2024.100081.

2. Santos-Júnior, C.D.; Torres, M.D.T.; Duan, Y.; Rodríguez del Río, Á.; Schmidt, T.S.B.; Chong, H.; Fullam, A.; Kuhn, M.; Zhu, C.; Houseman, A.; et al. Discovery of antimicrobial peptides in the global microbiome with machine learning. Cell 2024, 187, 3761–3778.e3716, 10.1016/j.cell.2024.05.013.

3. Purohit, K.; Reddy, N.; Sunna, A. Exploring the Potential of Bioactive Peptides: From Natural Sources to Therapeutics. Int J Mol Sci 2024, 25, doi:10.3390/ijms25031391.

4. Tucker, A.T.; Leonard, S.P.; DuBois, C.D.; Knauf, G.A.; Cunningham, A.L.; Wilke, C.O.; Trent, M.S.; Davies, B.W. Discovery of Next-Generation Antimicrobials through Bacterial Self-Screening of Surface-Displayed Peptide Libraries. Cell 2018, 172, 618–628.e613, doi:10.1016/j.cell.2017.12.009.

5. Mendes, B.; Edwards-Gayle, C.; Barrett, G. Peptide lipidation and shortening optimises antibacterial, antibiofilm and membranolytic actions of an amphiphilic polylysine-polyphenyalanine octapeptide. Current Research in Biotechnology 2024, 8, 100240, 10.1016/j.crbiot.2024.100240.

6. Mendes, B.; Almeida, J.R.; Vale, N.; Gomes, P.; Gadelha, F.R.; Da Silva, S.L.; Miguel, D.C. Potential use of 13-mer peptides based on phospholipase and oligoarginine as leishmanicidal agents. Comparative Biochemistry and Physiology Part C: Toxicology & Pharmacology 2019, 226, 108612, 10.1016/j.cbpc.2019.108612.

7. Duca, S.; Nikoi, N.D.; Berrow, M.; Barber, L.; Slope, L.N.; Peacock, A.F.A.; de Cogan, F. Oligoarginine peptide structure and its effect on cell penetration in ocular drug delivery. Heliyon 2024, 10, e35109, 10.1016/j.heliyon.2024.e35109.

8. Gopal, R.; Kim, Y.J.; Seo, C.H.; Hahm, K.S.; Park, Y. Reversed sequence enhances antimicrobial activity of a synthetic peptide. J Pept Sci 2011, 17, 329–334, doi:10.1002/psc.1369.

9. Adak, A.; Castelletto, V.; de Mello, L.; Mendes, B.; Barrett, G.; Seitsonen, J.; Hamley, I.W. Effect of Chirality and Amphiphilicity on the Antimicrobial Activity of Tripodal Lysine-Based Peptides. ACS Applied Bio Materials 2025, 8, 803–813, doi:10.1021/acsabm.4c01635.

10. Edwards-Gayle, C.J.C.; Barrett, G.; Roy, S.; Castelletto, V.; Seitsonen, J.; Ruokolainen, J.; Hamley, I.W. Selective Antibacterial Activity and Lipid Membrane Interactions of Arginine-Rich Amphiphilic Peptides. ACS Applied Bio Materials 2020, 3, 1165–1175, doi:10.1021/acsabm.9b00894.

11. Zhang, R.; Yan, H.; Wang, X.; Cong, H.; Yu, B.; Shen, Y. Screening of a short chain antimicrobial peptide-FWKFK and its application in wound healing. Biomater Sci 2023, 11, 1867–1875, doi:10.1039/d2bm01992b.

12. Edwards-Gayle, C.J.C.; Castelletto, V.; Hamley, I.W.; Barrett, G.; Greco, F.; Hermida-Merino, D.; Rambo, R.P.; Seitsonen, J.; Ruokolainen, J. Self-Assembly, Antimicrobial Activity, and Membrane Interactions of Arginine-Capped Peptide Bola-Amphiphiles. ACS Applied Bio Materials 2019, 2, 2208–2218, doi:10.1021/acsabm.9b00172.

13. Castelletto, V.; Barnes, R.H.; Karatzas, K.-A.; Edwards-Gayle, C.J.C.; Greco, F.; Hamley, I.W.; Rambo, R.; Seitsonen, J.; Ruokolainen, J. Arginine-Containing Surfactant-Like Peptides: Interaction with Lipid Membranes and Antimicrobial Activity. Biomacromolecules 2018, 19, 2782–2794, doi:10.1021/acs.biomac.8b00391.

14. Luo, X.; Ye, X.; Ding, L.; Zhu, W.; Yi, P.; Zhao, Z.; Gao, H.; Shu, Z.; Li, S.; Sang, M.;, et al. Fine-Tuning of Alkaline Residues on the Hydrophilic Face Provides a Non-toxic Cationic α-Helical Antimicrobial Peptide Against Antibiotic-Resistant ESKAPE Pathogens. Frontiers in Microbiology 2021, 12, doi:10.3389/fmicb.2021.684591.

15. Bacalum, M.; Radu, M. Cationic Antimicrobial Peptides Cytotoxicity on Mammalian Cells: An Analysis Using Therapeutic Index Integrative Concept. International Journal of Peptide Research and Therapeutics 2015, 21, 47–55, doi:10.1007/s10989-014-9430-z.

16. Almeida, J.R.; Palacios, A.L.V.; Patiño, R.S.P.; Mendes, B.; Teixeira, C.A.S.; Gomes, P.; da Silva, S.L. Harnessing snake venom phospholipases A(2) to novel approaches for overcoming antibiotic resistance. Drug Dev Res 2019, 80, 68–85, doi:10.1002/ddr.21456.

17. Rounds, T.; Straus, S.K. Lipidation of Antimicrobial Peptides as a Design Strategy for Future Alternatives to Antibiotics. Int J Mol Sci 2020, 21, doi:10.3390/ijms21249692.

18. Li, Y.; Clark, K.A.; Tan, Z. Methods for engineering therapeutic peptides. Chinese Chemical Letters 2018, 29, 1074–1078, 10.1016/j.cclet.2018.05.027.

19. Meena, K.R.; Kanwar, S.S. Lipopeptides as the antifungal and antibacterial agents: applications in food safety and therapeutics. Biomed Res Int 2015, 2015, 473050, doi:10.1155/2015/473050.

20. Steenbergen, J.N.; Alder, J.; Thorne, G.M.; Tally, F.P. Daptomycin: a lipopeptide antibiotic for the treatment of serious Gram-positive infections. Journal of Antimicrobial Chemotherapy 2005, 55, 283–288, doi:10.1093/jac/dkh546.

21. Schneider, T.; Müller, A.; Miess, H.; Gross, H. Cyclic lipopeptides as antibacterial agents - potent antibiotic activity mediated by intriguing mode of actions. Int J Med Microbiol 2014, 304, 37–43, doi:10.1016/j.ijmm.2013.08.009.

22. Wang, J.; Sintim, H.O. Antibiotics That Disrupt Cell Wall and Bacterial Membrane Formation and Integrity. In Reference Module in Biomedical Sciences; Elsevier: 2014.

23. Decker, A.P.; Mechesso, A.F.; Wang, G. Expanding the Landscape of Amino Acid-Rich Antimicrobial Peptides: Definition, Deployment in Nature, Implications for Peptide Design and Therapeutic Potential. Int J Mol Sci 2022, 23, doi:10.3390/ijms232112874.

24. Fan, L.; Sun, J.; Zhou, M.; Zhou, J.; Lao, X.; Zheng, H.; Xu, H. DRAMP: a comprehensive data repository of antimicrobial peptides. Scientific Reports 2016, 6, 24482, doi:10.1038/srep24482.

25. O’Toole, G.A. Microtiter dish biofilm formation assay. Journal of visualized experiments: JoVE 2011, 2437.

26. Peña-Carrillo, M.S.; Pinos-Tamayo, E.A.; Mendes, B.; Domínguez-Borbor, C.; Proaño-Bolaños, C.; Miguel, D.C.; Almeida, J.R. Dissection of phospholipases A(2) reveals multifaceted peptides targeting cancer cells, Leishmania and bacteria. Bioorg Chem 2021, 114, 105041, doi:10.1016/j.bioorg.2021.105041.

27. Almeida, J.R. The Century-Long Journey of Peptide-Based Drugs. Antibiotics 2024, 13, 196.

28. Cresti, L.; Cappello, G.; Pini, A. Antimicrobial Peptides towards Clinical Application-A Long History to Be Concluded. Int J Mol Sci 2024, 25, doi:10.3390/ijms25094870.

29. Mookherjee, N.; Anderson, M.A.; Haagsman, H.P.; Davidson, D.J. Antimicrobial host defence peptides: functions and clinical potential. Nature Reviews Drug Discovery 2020, 19, 311–332, doi:10.1038/s41573-019-0058-8.

30. Botelho Sampaio de Oliveira, K.; Lopes Leite, M.; Albuquerque Cunha, V.; Brito da Cunha, N.; Luiz Franco, O. Challenges and advances in antimicrobial peptide development. Drug Discovery Today 2023, 28, 103629, 10.1016/j.drudis.2023.103629.

31. Xuan, J.; Feng, W.; Wang, J.; Wang, R.; Zhang, B.; Bo, L.; Chen, Z.-S.; Yang, H.; Sun, L. Antimicrobial peptides for combating drug-resistant bacterial infections. Drug Resistance Updates 2023, 68, 100954, 10.1016/j.drup.2023.100954.

32. Li, J.; Koh, J.-J.; Liu, S.; Lakshminarayanan, R.; Verma, C.S.; Beuerman, R.W. Membrane Active Antimicrobial Peptides: Translating Mechanistic Insights to Design. Frontiers in Neuroscience 2017, 11, doi:10.3389/fnins.2017.00073.

33. Zhu, M.; Chen, J.; Lin, Y. Exploring chemical space and structural diversity of supramolecular peptide materials. Supramolecular Materials 2023, 2, 100030, 10.1016/j.supmat.2022.100030.

34. Capecchi, A.; Reymond, J.-L. Peptides in chemical space. Medicine in Drug Discovery 2021, 9, 100081, 10.1016/j.medidd.2021.100081.

35. Yang, B.; Yang, H.; Liang, J.; Chen, J.; Wang, C.; Wang, Y.; Wang, J.; Luo, W.; Deng, T.; Guo, J. A review on the screening methods for the discovery of natural antimicrobial peptides. Journal of Pharmaceutical Analysis 2025, 15, 101046, 10.1016/j.jpha.2024.101046.

36. Szymczak, P.; Szczurek, E. Artificial intelligence-driven antimicrobial peptide discovery. Current Opinion in Structural Biology 2023, 83, 102733, 10.1016/j.sbi.2023.102733.

37. Chang, L.; Mondal, A.; Singh, B.; Martínez-Noa, Y.; Perez, A. Revolutionizing peptide-based drug discovery: Advances in the post-AlphaFold era. WIREs Computational Molecular Science 2024, 14, e1693, 10.1002/wcms.1693.

38. Almeida, J.R.; Mendes, B.; Lancellotti, M.; Franchi, G.C.; Passos, Ó.; Ramos, M.J.; Fernandes, P.A.; Alves, C.; Vale, N.; Gomes, P.;, et al. Lessons from a Single Amino Acid Substitution: Anticancer and Antibacterial Properties of Two Phospholipase A2-Derived Peptides. Current Issues in Molecular Biology 2022, 44, 46–62.

39. Feijoo-Coronel, M.L.; Mendes, B.; Ramírez, D.; Peña-Varas, C.; de los Monteros-Silva, N.Q.E.; Proaño-Bolaños, C.; de Oliveira, L.C.; Lívio, D.F.; da Silva, J.A.; da Silva, J.M.S.F.; et al. Antibacterial and Antiviral Properties of Chenopodin-Derived Synthetic Peptides. Antibiotics 2024, 13, 78.

40. Bello-Madruga, R.; Torrent Burgas, M. The limits of prediction: Why intrinsically disordered regions challenge our understanding of antimicrobial peptides. Computational and Structural Biotechnology Journal 2024, 23, 972–981, 10.1016/j.csbj.2024.02.008.

41. Cardoso, M.H.; Orozco, R.Q.; Rezende, S.B.; Rodrigues, G.; Oshiro, K.G.N.; Cândido, E.S.; Franco, O.L. Computer-Aided Design of Antimicrobial Peptides: Are We Generating Effective Drug Candidates? Frontiers in Microbiology 2020, 10, doi:10.3389/fmicb.2019.03097.

42. Joondan, N.; Jhaumeer-Laulloo, S.; Caumul, P. A study of the antibacterial activity of l-Phenylalanine and l-Tyrosine esters in relation to their CMCs and their interactions with 1,2-dipalmitoyl-sn-glycero-3-phosphocholine, DPPC as model membrane. Microbiological Research 2014, 169, 675–685, 10.1016/j.micres.2014.02.010.

43. Valdivieso-Rivera, F.; Bermúdez-Puga, S.; Proaño-Bolaños, C.; Almeida, J.R. Deciphering the Limitations and Antibacterial Mechanism of Cruzioseptins. International Journal of Peptide Research and Therapeutics 2022, 28, 73, doi:10.1007/s10989-022-10383-4.

44. Lu, J.; Xu, H.; Xia, J.; Ma, J.; Xu, J.; Li, Y.; Feng, J. D- and Unnatural Amino Acid Substituted Antimicrobial Peptides With Improved Proteolytic Resistance and Their Proteolytic Degradation Characteristics. Frontiers in Microbiology 2020, 11, doi:10.3389/fmicb.2020.563030.

45. Lin, B.; Hung, A.; Singleton, W.; Darmawan, K.K.; Moses, R.; Yao, B.; Wu, H.; Barlow, A.; Sani, M.-A.; Sloan, A.J.;, et al. The effect of tailing lipidation on the bioactivity of antimicrobial peptides and their aggregation tendency. Aggregate 2023, 4, e329, 10.1002/agt2.329.

46. Makowska, M.; Wardowska, A.; Bauer, M.; Wyrzykowski, D.; Małuch, I.; Sikorska, E. Impact of lipidation site on the activity of α-helical antimicrobial peptides. Bioorganic Chemistry 2024, 153, 107821, 10.1016/j.bioorg.2024.107821.

47. Grimsey, E.; Collis, D.W.P.; Mikut, R.; Hilpert, K. The effect of lipidation and glycosylation on short cationic antimicrobial peptides. Biochimica et Biophysica Acta (BBA) - Biomembranes 2020, 1862, 183195, 10.1016/j.bbamem.2020.183195.

48. Saghiri, A.; Reza Bozorgmehr, M.; Morsali, A. Deciphering the Impact of Cyclization and Lysine Charges on Antimicrobial Peptides Using Molecular Dynamics Simulations and Density Functional Theory. ChemistrySelect 2024, 9, e202401879, 10.1002/slct.202401879.

49. Garton, M.; Nim, S.; Stone, T.A.; Wang, K.E.; Deber, C.M.; Kim, P.M. Method to generate highly stable D-amino acid analogs of bioactive helical peptides using a mirror image of the entire PDB. Proceedings of the National Academy of Sciences 2018, 115, 1505–1510, doi:doi:10.1073/pnas.1711837115.

50. Upert, G.; Luther, A.; Obrecht, D.; Ermert, P. Emerging peptide antibiotics with therapeutic potential. Medicine in Drug Discovery 2021, 9, 100078, 10.1016/j.medidd.2020.100078.

51. Kang, S.-J.; Nam, S.H.; Lee, B.-J. Engineering Approaches for the Development of Antimicrobial Peptide-Based Antibiotics. Antibiotics 2022, 11, doi:10.3390/antibiotics11101338.

52. Zhang, L.; Rozek, A.; Hancock, R.E.W. Interaction of Cationic Antimicrobial Peptides with Model Membranes*. Journal of Biological Chemistry 2001, 276, 35714–35722, 10.1074/jbc.M104925200.

53. Wimley, W.C. Describing the mechanism of antimicrobial peptide action with the interfacial activity model. ACS Chem Biol 2010, 5, 905–917, doi:10.1021/cb1001558.

54. Oyinloye, B.E.; Adenowo, A.F.; Kappo, A.P. Reactive oxygen species, apoptosis, antimicrobial peptides and human inflammatory diseases. Pharmaceuticals (Basel) 2015, 8, 151–175, doi:10.3390/ph8020151.

55. Rowe-Magnus, D.A.; Kao, A.Y.; Prieto, A.C.; Pu, M.; Kao, C. Cathelicidin Peptides Restrict Bacterial Growth via Membrane Perturbation and Induction of Reactive Oxygen Species. mBio 2019, 10, doi:10.1128/mBio.02021-19.

56. Schmidt, N.; Mishra, A.; Lai, G.H.; Wong, G.C.L. Arginine-rich cell-penetrating peptides. FEBS Letters 2010, 584, 1806–1813, 10.1016/j.febslet.2009.11.046.

57. Sun, A.; Huang, Z.; He, L.; Dong, W.; Tian, Y.; Huang, A.; Wang, X. Metabolomic analyses reveal the antibacterial properties of a novel antimicrobial peptide MOp3 from Moringa oleifera seeds against Staphylococcus aureus and its application in the infecting pasteurized milk. Food Control 2023, 150, 109779, 10.1016/j.foodcont.2023.109779.

58. Bermúdez-Puga, S.; Dias, M.; Freire de Oliveira, T.; Mendonça, C.M.N.; Yokomizo de Almeida, S.R.; Rozas, E.E.; do Nascimento, C.A.O.; Mendes, M.A.; Oliveira De Souza de Azevedo, P.; Almeida, J.R.;, et al. Dual antibacterial mechanism of [K4K15]CZS-1 against Salmonella Typhimurium: a membrane active and intracellular-targeting antimicrobial peptide. Front Microbiol 2023, 14, 1320154, doi:10.3389/fmicb.2023.1320154.

59. Ditsawanon, T.; Phaonakrob, N.; Roytrakul, S. Mechanisms of Antimicrobial Peptides from Bagasse against Human Pathogenic Bacteria. Antibiotics 2023, 12, doi:10.3390/antibiotics12030448.

60. Tomasinsig, L.; Scocchi, M.; Mettulio, R.; Zanetti, M. Genome-wide transcriptional profiling of the Escherichia coli response to a proline-rich antimicrobial peptide. Antimicrob Agents Chemother 2004, 48, 3260–3267, doi:10.1128/aac.48.9.3260-3267.2004.

61. Georgieva, M.; Heinonen, T.; Vitale, A.; Hargraves, S.; Causevic, S.; Pillonel, T.; Eberl, L.; Widmann, C.; Jacquier, N. Bacterial surface properties influence the activity of the TAT-RasGAP317-326 antimicrobial peptide. iScience 2021, 24, 102923, 10.1016/j.isci.2021.102923.

62. Hong, R.W.; Shchepetov, M.; Weiser, J.N.; Axelsen, P.H. Transcriptional profile of the Escherichia coli response to the antimicrobial insect peptide cecropin A. Antimicrob Agents Chemother 2003, 47, 1–6, doi:10.1128/aac.47.1.1-6.2003.

63. Tang, Y.; Yang, C.; Zhao, J.; Heng, H.; Peng, M.; Sun, L.; Dai, L.; Chan, E.W.- C.; Chen, S. LTX-315 is a novel broad-spectrum antimicrobial peptide against clinical multidrug-resistant bacteria. Journal of Advanced Research 2025, 10.1016/j.jare.2024.12.044.

64. Cardoso, M.H.; de Almeida, K.C.; Cândido, E.S.; Fernandes, G.d.R.; Dias, S.C.; de Alencar, S.A.; Franco, O.L. Comparative transcriptome analyses of magainin I-susceptible and -resistant Escherichia coli strains. Microbiology 2018, 164, 1383–1393, 10.1099/mic.0.000725.

65. Zhao, K.; Liu, M.; Burgess, R.R. Adaptation in bacterial flagellar and motility systems: from regulon members to ‘foraging’-like behavior in E. coli. Nucleic Acids Research 2007, 35, 4441–4452, doi:10.1093/nar/gkm456.

66. Ikeda, T.; Shinagawa, T.; Ito, T.; Ohno, Y.; Kubo, A.; Nishi, J.; Gotoh, Y.; Ogura, Y.; Ooka, T.; Hayashi, T. Hypoosmotic stress induces flagellar biosynthesis and swimming motility in Escherichia albertii. Communications Biology 2020, 3, 87, doi:10.1038/s42003-020-0816-5.

67. Avci, F.G.; Akbulut, B.S.; Ozkirimli, E. Membrane Active Peptides and Their Biophysical Characterization. Biomolecules 2018, 8, doi:10.3390/biom8030077.

68. Andrews, S.C.; Robinson, A.K.; Rodríguez-Quiñones, F. Bacterial iron homeostasis. FEMS Microbiology Reviews 2003, 27, 215–237, doi:10.1016/s0168-6445(03)00055-x.

69. Tawfik Mohamed, M.; Bertelsen, M.; Abdel-Rahman Mohamed, A.; Strong Peter, N.; Miller, K. Scorpion Venom Antimicrobial Peptides Induce Siderophore Biosynthesis and Oxidative Stress Responses in Escherichia coli. mSphere 2021, *6*, 10.1128/msphere.00267-00221, doi:10.1128/msphere.00267-21.

70. Pletzer, D.; Mansour, S.C.; Hancock, R.E.W. Synergy between conventional antibiotics and anti-biofilm peptides in a murine, sub-cutaneous abscess model caused by recalcitrant ESKAPE pathogens. PLOS Pathogens 2018, 14, e1007084, doi:10.1371/journal.ppat.1007084.

71. Duan, H.; Zhang, X.; Li, Z.; Yuan, J.; Shen, F.; Zhang, S. Synergistic effect and antibiofilm activity of an antimicrobial peptide with traditional antibiotics against multi-drug resistant bacteria. Microbial Pathogenesis 2021, 158, 105056, 10.1016/j.micpath.2021.105056.

72. de la Fuente-Núñez, C.; Cardoso, M.H.; de Souza Cândido, E.; Franco, O.L.; Hancock, R.E.W. Synthetic antibiofilm peptides. Biochimica et Biophysica Acta (BBA) - Biomembranes 2016, 1858, 1061–1069, 10.1016/j.bbamem.2015.12.015.

